# Microbial functional guilds and genes are key to explaining soil nutrient cycling alongside soil and plant variables

**DOI:** 10.1101/2024.12.13.627780

**Authors:** Corinne Vietorisz, Nahuel Policelli, Abigail Li, Lindsey A. Adams, Jennifer M. Bhatnagar

## Abstract

Microbes play central roles in soil nutrient cycling, yet a limited range of microbial community characteristics have been used to explain ecosystem nutrient cycling rates and their importance relative to plant and abiotic factors remains unclear. In this study, we assessed which of 126 commonly measured soil fungal and bacterial community characteristics best explained soil nitrogen (N) and phosphorus (P) cycling rates in temperate forests in the Northeastern U.S., as well as the relative contributions of microbial, plant, and abiotic factors. Using boosted regression tree modeling, we identified the microbial variables with the highest contributions to models explaining nutrient cycling rates: the relative abundances of ectomycorrhizal fungi and N-decomposition genes from oligotrophic bacteria were the most important for net ammonification, the relative abundances of indicator taxa in bacterial networks, nitrifying bacteria, and copiotrophic bacteria were the most important for net nitrification, and the relative abundance of fungal P-cycling oxidoreductase genes was the most important for net soil phosphate change. Microbial variables explained more variation than plant and abiotic variables in multivariate linear models of net nitrification and net phosphate release rates, but not net ammonification rates, which were largely explained by soil edaphic factors. Leaf litter traits were also important in explaining variation in net nitrification rates, and soil temperature was important in explaining rates of net phosphate release in soil. Collectively, our findings suggest that the N-cycling capacity of microbial functional guilds and fungal community P-cycling capacity should be incorporated into ecosystem biogeochemical models to improve our predictions and understanding of nutrient cycling and related ecological processes.

## INTRODUCTION

Soil microbes, including fungi and bacteria, are the driving force behind nutrient cycling within forest soils, secreting enzymes that release key elements like nitrogen (N) and phosphorus (P) from dead organic matter, immobilizing them in microbial biomass, and transferring these elements to plant hosts (Smith and Read 2008, Schneider et al. 2012). Despite their central functions in nutrient cycling, we still don’t understand which aspects of microbial communities best explain variation in soil nutrient cycling rates. Meta-analyses show that, on average, abiotic drivers alone such as climate, pH, and soil organic matter (SOM) can explain roughly half of the variation in nutrient cycling (Graham et al. 2016), while plant factors such as leaf litter quality and fine root distribution can explain between one-third to three-quarters of variation on their own (Scott and Binkley 1997, Thomas and Prescott 2000, Hobbie et al. 2015, Finzi et al. 2015). Incorporating microbial factors sometimes improves models explaining biogeochemical cycling, though bacterial community structure and diversity are often the only microbial factors considered (Graham et al. 2016). A wide range of soil microbial community characteristics can now be measured efficiently and at low cost, making them potential candidates to improve biogeochemical model predictions and our understanding of nutrient cycling and related ecological processes – such as carbon (C) cycling and primary productivity – across spatial and temporal scales.

Some microbial variables have been linked to nutrient cycling rates, such as microbial (usually bacterial) diversity, which can be positively correlated with ecosystem processes like N cycling, especially in species-poor communities (Salonius 1981, Setälä and McLean 2004, Bell et al. 2005, Philippot et al. 2013). The relative abundances of microbial functional guilds involved in nutrient cycling have also correlated with N cycling: soil nitrification rates can positively correlate with the relative abundance of ammonia-oxidizing prokaryotes (Isobe et al. 2015, Zhang et al. 2019, Sorensen et al. 2019, Tatsumi et al. 2020) and net N mineralization rates can positively correlate with the relative abundance of ectomycorrhizal fungi (EMF, Sorensen et al. 2019, Zhang et al. 2023). Additionally, abundances of functional marker genes encoding enzymes that perform nutrient transformations (primarily bacterial genes) have sometimes correlated with N cycling processes (Petersen et al. 2012, Isobe et al. 2015, Tang et al. 2019, Chen et al. 2020), but other studies find no correlation (Philippot et al. 2013, Graham et al. 2014, Rocca et al. 2015, Zhang et al. 2021, Saifuddin et al. 2021). This discrepancy could be because community-level functional gene abundances are not linked to microbial taxonomy, marker genes may not be representative of a complete metabolic pathway (Albright et al. 2019), or gene copy abundances may not reflect gene or protein expression levels in the community.

Comprehensive groups of genes involved in biochemical transformation pathways from relevant microbial functional guilds, rather than total abundances of a single gene, may be a better predictor of nutrient cycling processes, as nutrient cycling gene abundances and expression can be related to a microbe’s functional guild (Talbot et al. 2015, Romero-Olivares et al. 2021, Auer and Buée et al. 2024). For example, ectomycorrhizal fungi (EMF) can have higher gene expression of chitinases and other fungal cell wall-degrading enzymes than saprotrophic fungi (Maillard et al. 2022, Auer and Buée et al. 2024), so chitinase genes from EMF may best explain rates of chitin degradation in soils. P decomposition rates in soils may also be explained by organic matter-degrading genes from mycorrhizal fungi, as they are hypothesized to perform decomposition to acquire nutrients instead of C (Talbot et al. 2015) and many EMF have diverse methods of P-mobilization in soils (Plassard et al. 2011). Soil nitrification rates may be best explained by nitrification gene abundances from narrow groups of nitrifying bacteria. For example, nitrification rates in wastewater correlate with nitrification gene expression in communities dominated by a single order of bacteria (*Nitrosomonodales*, Kapoor et al. 2016). The relative abundances of functional gene copies derived from specific microbial guilds has been shown to correlate with tree growth (Anthony et al. 2022), but remains untested for soil nutrient cycling.

Another untested microbial metric that may explain soil nutrient cycling rates is associations between microbial groups, as interactions between species can drive community assembly and function (Guidi et al. 2016, Maynard et al. 2018, Ratzke et al. 2020). For example, competition for ammonium between EMF and ammonia-oxidizing prokaryotes may decrease nitrification rates (Tatsumi et al. 2020), so the correlation between these two groups could explain variation in net nitrification rates. Networks of commonly co-occurring taxa have also been linked to certain ecosystem processes (Guidi et al. 2016, Wagg et al. 2019) and abundances of indicator taxa central to these networks have the potential to explain nutrient cycling rates. Despite the diversity of microbial community metrics that are relatively easy to measure, it is unknown which of these metrics best explain soil N and P cycling rates, or their importance relative to plant and soil abiotic factors.

In this study, we sought to determine which commonly measured soil microbial community characteristics best explain soil N and P cycling and the relative contributions of microbial, plant, and abiotic factors in explaining these processes. We hypothesized that: 1) relative gene abundances from specific functional guilds will be the best microbial metric to explain rates of inorganic soil N and P cycling. We expected that net ammonification and net phosphate change would be best explained by EMF N- and P-decomposition gene abundances, respectively, while net nitrification rates would be best explained by nitrifying bacterial nitrification gene abundances; and 2) microbial metrics will explain more of the variation in models explaining soil N and P cycling rates than plant or edaphic factors. To test our hypotheses, we sampled soils across six mixed deciduous mid-successional temperate forest sites in Massachusetts, USA that maximized variation in soil microbial communities, vegetation, and soil edaphic factors. We measured soil N and P cycling rates, fungal and bacterial community characteristics, plant community characteristics, and soil abiotic attributes. We assessed the relative contributions of microbial, plant, and abiotic factors in explaining rates of soil N and P cycling using a combination of boosted regression trees (BRTs) and multivariate linear models. We considered the “best” variables explaining nutrient cycling rates to be those that explained the most variation in N and P cycling rates within the models.

## METHODS

### Site description and sampling design

We sampled a set of temperate forest sites in eastern and central Massachusetts, USA that maximized variation in soil microbial community composition, plant community characteristics, and soil nutrient cycling rates, but minimized confounding correlations between microbial and plant factors. We selected six field sites (Appendix S1: Section S1: Methods), where each site contains 4 forest stand types, all within 2.5 km of each other: mature white pine (*Pinus strobus*)- dominated stands, mature hardwood-dominated stands (mostly *Quercus* spp.), hardwood-dominated stands with white pine saplings encroaching into the understory, and mixed mature white pine and hardwood stands (Appendix S1: Fig. S1). The stands with pine encroachment contained no mature pine trees, but had scattered white pine saplings throughout the understory. Young pines associate with unique pine-specific EMF (i.e., EMF that form host-specific associations with pines, Policelli et al. 2020), but contribute very little pine litter to the forest floor (Appendix S1: Fig. S2), allowing us to isolate effects of pine-associated soil microbial communities from litter inputs on soil nutrient cycling. Within each forest type at each site, one transect was laid from the forest edge to interior, perpendicular to the forest edge (6 sites x 4 forest types = 24 total transects), as shown in Appendix S1: Fig. S1. We included a forest edge to interior transect because microbial communities and soil characteristics vary with distance from the forest edge in these forests, even when plant communities do not (Tatsumi et al. 2023), further isolating the effects of plant and microbial communities on soil nutrient cycling. Two sampling points were established within 5 m from the transect at 0 m, 15 m, 30 m, 60 m, and 90 m from the forest edge (Appendix S1: Fig. S1). This resulted in a total of 204 sampling points (24 transects x 4 or 5 distances per transect x 2 sampling points per distance).

### Soil sampling, nutrient cycling and edaphic factors

The top 6 cm of soil (primarily organic horizon) was sampled for microbial metrics, nitrogen cycling rates, and edaphic measurements in July 2021, and for phosphorus cycling rates in July 2022 as described in Appendix S1: Section S1: Methods. During the bulk soil collections, fresh sieved soil was saved at 4℃ for inorganic nutrient extractions, pH, and moisture measurements, an aliquot of sieved soil was frozen at -80℃ for subsequent DNA extraction, and an aliquot of sieved soil was air-dried for total elemental analysis. During soil sampling in July 2021 and 2022, soil temperature (Rapitest digital soil thermometer, Luster Leaf, Woodstock, IL) and depth of organic horizon were measured directly along the transect, equidistant between each A and B sampling point at all distances. In the same 204 soil samples from July 2021 that were used for inorganic N extractions, % soil organic matter, gravimetric soil moisture, and pH were measured following the protocols described in Caron et al. (2023).

To measure soil net ammonification and nitrification, we used the “buried bag” method, where an initial soil sample is taken to measure ammonium and nitrate, then another soil sample is buried beneath the litter layer in the same location within a polyethylene bag and incubated in the field for a month (Eno 1960, Westermann & Crothers 1980, Hanselman et al. 2004, Durán et al. 2012, Caron et al. 2023). Initial soil sampling was conducted in mid-July 2021, and the buried bags were collected 4 weeks later in mid-August 2021. Soil samples were collected, extracted in 2M potassium chloride, and analyzed colorimetrically for inorganic nitrogen (ammonium and nitrate) content as described in Appendix S1: Section S1: Methods. To measure net soil phosphate change, a second year of soil sampling was completed in early July 2022 – early August 2022. We followed the same buried bag and soil sampling protocol in the same sampling locations as for the net ammonification and net nitrification assays, except that we measured Olsen inorganic P (a measure of plant-available P, Olsen et al. 1954). We performed extractions in 0.5M sodium bicarbonate and analyzed the extracts for phosphate concentrations colorimetrically as described in Appendix S1: Section S1: Methods (Olsen et al. 1954, Frank et al. 1998). On the 144 soils from the 0 m, 15 m, and 60 m distances collected in 2021, an aliquot of sieved soil was air dried and sent to the Ohio State Service Testing and Research Laboratory for total phosphorus analysis via the EPA 3051A acid digestion (U.S. EPA 2007). The same 144 soils were also dried at 65℃, ground to a fine powder in a ball-mill, and combusted on an elemental analyzer (Vario EL cube, Elementar) for total C and N analysis.

### Vegetation factors

In September 2021, forest floor leaf litter depth and composition, and root density were measured at each distance from the forest edge along the transect as described in Appendix S1: Section S1: Methods. In September 2021 – September 2022, litterfall weight and chemistry were measured once per transect as described in Appendix S1: Section S1: Methods. To measure aboveground vegetation community composition, in June 2022, a 10m x 10m square plot at each transect was created at each distance from the forest edge centered around the sampling points. Within the plot, we measured the total basal area of canopy trees by species and understory plant cover by growth form using the Braun-Blanquet method (Braun-Blanquet 1932, Matteucci and Colma 1982) as described in Appendix S1: Section S1: Methods.

### Microbial DNA extraction, amplicon sequencing, bioinformatics, and qPCR

Total soil DNA was extracted from approximately 0.25 g of each soil sample (204 samples) using the DNeasy PowerSoil Kit (QIAGEN, Hilden, Germany). To amplify microbial DNA, we used modified versions of the primer set fITS7 and ITS4 for fungi (amplifying the ITS2 region of rDNA, Anthony et al. 2017) and modified versions of the primer set 515f and 806r for bacteria (amplifying the v4 16S region of rDNA, Caporaso et al. 2011) that contained both the Illumina adapter and individual sample indexes. DNA amplification and sequencing was performed following the procedure outlined by Tatsumi et al (2023) and in Appendix S1: Section S1: Methods. Bioinformatics were performed in R (version 4.2.1), where the R package dada2 was used for sequence quality control, paired-end assembly, identification of amplicon sequence variants (ASVs), and taxonomy assignment (Caporaso et al. 2011, Callahan et al. 2016), with any deviations from default noted in Appendix S1: Section S1: Methods. To identify and remove outliers based on community composition and sequencing depth, we ran a Non-Metric Multidimensional Scaling analysis using an Aitchison distance matrix (Gloor et al. 2017, Appendix S1: Section S1: Methods). 13 samples with under 8000 reads were removed for ITS sequences, and 3 samples determined as outliers due to sequencing errors or contamination were removed for 16S sequences.

To estimate total fungal and bacterial abundances, total ITS2 and 16S v4 gene abundances were quantified on all DNA extracts via qPCR using the same primer sets used for amplicon sequencing (without the Illumina adapter and indexes, White et al. 1990, Caporaso et al. 2011), following the protocol outlined in Tatsumi et al. (2023).

### Microbial community metrics

Based on DNA sequence analysis, we quantified 126 microbial community characteristics across 7 different categories of microbial community traits (Table 1). To obtain the relative abundances of fungal and bacterial functional guilds, the ITS and 16S ASV tables were converted to relative abundances by dividing each ASV count in each sample by the total number of reads in that sample (Weiss et al. 2017). Fungal genera were assigned functional guilds and ectomycorrhizal fungi were assigned exploration types using the FungalTraits database (Põlme et al. 2020). Bacterial taxa were assigned as copiotrophs or oligotrophs, and/or nitrifying bacteria, using a database compiled from literature reviews and genomic pathway presence where we assigned guilds at the genus through phylum level (Ho et al. 2017, Albright et al. 2019, Naylor et al. 2020, Averill et al. 2021), which allowed us to include many bacterial ASVs that did not have taxonomy assignments at lower taxonomic levels. The relative abundance of functional guilds in each sample was calculated by summing the relative abundances of ASVs belonging to each functional guild. Fungal and bacterial alpha diversity were calculated on rarefied ASV tables (Weiss et al. 2017) using Shannon’s diversity index. 16S samples were rarefied to a depth of 14,906 reads, and ITS samples to a depth of 7,345 reads. Fungal and bacterial evenness were calculated by dividing the Shannon’s diversity index by the natural log of the richness (Pielou 1966).

**Table 1.** Definitions of the microbial community characteristics used to explain nutrient cycling rates.

| Microbial community characteristic type | Definition | Example characteristic |
| --- | --- | --- |
| Functional guild relative abundances | The summed relative abundance of ASVs belonging to a microbial functional group within a sample | Relative abundance of ectomycorrhizal fungi |
| Functional gene relative abundances | The estimated relative abundance of functional gene copies in the bacterial or fungal communities in a sample; calculated via PICRUST2 (for bacteria) or using fungal genome annotations (for fungi) in a sample | Relative abundance of fungal chitinase genes |
| Functional gene relative abundances within functional guilds | The estimated relative abundance of functional gene copies, summed for a functional guild in a sample | Relative abundance of chitinase genes from ectomycorrhizal fungi |
| Indicator co-occurrence modules (ICMs) | The summed relative abundance of all fungal or bacterial ASVs in an indicator co-occurrence module (i.e. group of co-occurring ASVs identified via network analysis) in a sample. | Relative abundance of the fungal “high ammonification” ICM |
| Diversity metrics | Metrics of fungal or bacterial alpha diversity | Fungal Shannon diversity |
| 16S and ITS gene abundances | The total number of 16S v4 or ITS2 gene copies in a sample (per µl DNA extract) as determined by qPCR; a proxy for total bacterial and fungal abundance, respectively | Fungal ITS gene copies |
| Functional guild correlations | The strength of the Spearman correlation between the proportion of a bacterial and fungal functional guild, calculated at the transect-level using all samples within a transect | Correlation between ectomycorrhizal fungi and nitrifying bacteria |

To estimate the relative abundances of functional genes, Enzyme Commission (E.C.) numbers per ASV per sample were estimated via PICRUSt2 for bacteria (Douglas et al. 2020), and via the method described in Anthony et al (2022) with modifications by Atherton et al. (in preparation) for fungi. To calculate the relative abundances of genes encoding enzymes involved in N and P cycling in each sample, we summed the relative abundances of all E.C. numbers encoding the enzymes listed in Appendix S1: Table S1. We focused only on N- and P-cycling genes directly involved in ammonification, nitrification, and phosphate release from organic matter, which are well-annotated in environmental samples relative to other gene groups (Albright et al. 2019, Zeng et al. 2022). To calculate the relative abundances of genes derived from specific functional guilds in each sample, we summed the relative abundances of all E.C. numbers encoding N and P cycling enzymes that are derived from each functional guild.

In addition, we created “indicator co-occurrence modules” (ICMs) of fungal or bacterial taxa that are central to microbial networks associated with each nutrient cycling rate. We first ran a Weighted Gene Correlation Network Analysis (WGCNA) using the ‘WGCNA’ (Langfelder and Horvath 2008) and ‘flashClust’ (Langfelder and Horvath 2012) R packages on our 16S and ITS ASV tables as described in Appendix S1: Section S1: Methods. This analysis identifies microbial community network structures that are positively or negatively associated with nutrient cycling rates. We then identified the taxa that were most central to each network module (using a statistical metric of module membership, kME, Langfelder and Horvath 2008) and created a new metric of ICM relative abundance by summing their relative abundances. Taxa included in each module are shown in Appendix S1: Figures S3 and S4.

The strength of correlations between fungal and bacterial guilds were also calculated for all soil samples within each of the 24 transects using Spearman correlations. Guild correlations were calculated between all combinations of a fungal guild (e.g., EMF or saprotrophic fungi) and a bacterial guild (copiotrophic, oligotrophic, or nitrifying bacteria).

### Statistical methods

To test our hypotheses, we selected a unique set of potential explanatory variables for modeling each nutrient cycling process. A microbial variable was included in a model only if it directly related to a nutrient cycling mechanism (for example, P-cycling gene relative abundances were not included for N-cycling processes, and vice versa). All plant and soil abiotic variables were included as potential explanatory variables for all processes. This yielded 111 total variables for net ammonification (82 microbial, 20 plant, 9 soil), 68 variables for net nitrification (39 microbial, 20 plant, 9 soil), and 86 variables for net phosphate change (57 microbial, 20 plant, 9 soil). To examine the strength of correlation between explanatory variables and nutrient cycling processes, we ran Pearson correlations between all potential explanatory variables and the respective nutrient cycling rate. To reduce the total number of variables input to the multivariate model selection processes, for net ammonification and net nitrification models, we only input variables that had a significant correlation (p < 0.05) with the nutrient cycling rates. For net phosphate change, which had significant correlations with few variables, we input variables that had at least a marginally significant correlation (p < 0.1). See Appendix S1: Section S1: Methods for further detail on input variable selection for each nutrient cycling rate. This process yielded 17 input variables for net ammonification, 17 for net nitrification, and 6 for net phosphate change.

To compare the relative contributions of all microbial, plant, and soil abiotic variables explaining variation in net ammonification, net nitrification, and net phosphate change, we ran boosted regression trees (BRTs), which determine the relative contributions of many potential variables to explaining nutrient cycling rates and can fit complex, non-linear relationships, do not require data transformation, and can handle interaction effects and correlations (Appendix S1: Figure S6) between predictors (De’ath 2007, Elith et al. 2008, Allen et al. 2017). All BRTs were fit using the R package ‘dismo’ (Elith et al. 2008) and parameters were optimized following the procedure described in Appendix S1: Section S1: Methods. Because there is slight variation in the predictor variable contributions in each run, we ran 1,000 replicate model runs for each BRT.

To examine univariate linear relationships between nutrient cycling rates and explanatory variables identified as most important by the BRTs, we ran univariate linear mixed-effects models using the ‘lme4’ package in R (Bates et al. 2015), where sampling site was included as a random effect. If necessary, response and predictor variables were square-root transformed to meet linear model assumptions. For net nitrification, zero-inflated generalized linear mixed-models were run using the ‘glmmTMB’ package (Brooks et al. 2017) with the link set to “log”.

To determine which linear combinations of variables best explain variation in each nutrient cycling process, we constructed multivariate linear models for each nutrient cycling process and selected the best-fit model that explained the greatest amount of variation in nutrient cycling rate using the fewest possible independent variables. For net ammonification and net phosphate change, best-fit models were chosen by selecting the models with lowest Akaike Information Criteria (AIC) using the ‘leaps’ package testing for the Mallows’ C-p score of each candidate model, which maximizes variation explained in the data, and minimizes correlation between predictors and number of independent variables (Mallows 1973, Miller 2024). Site was included as a random effect unless it did not explain any variance in the model. To construct multivariate models explaining net nitrification, we performed stepwise model selection manually on zero-inflated generalized linear mixed models using the ‘glmmTMB’ command (Brooks et al. 2017) as described in Appendix S1: Section S1: Methods.

To test for the relationship between soil N and P cycling rates and litterfall variables and microbial guild correlations, transect-level multivariate linear models were constructed. Litterfall variables and microbial guild correlations were measured once per transect, and all other variables were averaged by transect to create one observation per transect (24 total). Model selection was performed as described above using the ‘leaps’ package. Net nitrification was log transformed (plus a pseudocount of 0.01) prior to input. All multivariate and univariate linear model assumptions were checked by examining plots of the residuals vs. fitted values, normal quantile-quantile plot, and covariance between predictors.

## RESULTS

In contrast to our first hypothesis, relative gene abundances derived from specific functional guilds were not the most important microbial metrics explaining any soil N or P cycling rates in the BRTs or multivariate linear models. Instead, functional guild abundances were the microbial factor most strongly tied to soil N cycling. For net ammonification, the microbial variable with the highest contribution to the BRTs was the relative abundance of EMF (Fig. 1a,b). The relative abundance of N-decomposition genes from oligotrophic bacteria (Fig 1a,c) had the next-highest contribution to the BRTs, followed by the fungal high ammonification ICM (Fig. 1a,d), which all had positive relationships with net ammonification. The N- decomposition gene groups that most strongly positively correlated with net ammonification rates were chitinases and N-cycling glycosidases (hydrolysing O- and S-glycosyl compounds, Fig. 1c, Appendix S1: Fig. S8). In the fungal high ammonification ICM, 4 of the genera in the module are EMF that occur frequently in our study sites, with these 4 EMF genera comprising 34% of all reads in our dataset. Ammonification also negatively correlated with fungal and bacterial diversity metrics, most strongly with bacterial Shannon diversity and fungal richness (Appendix S1: Fig. S8).

**Fig. 1.**
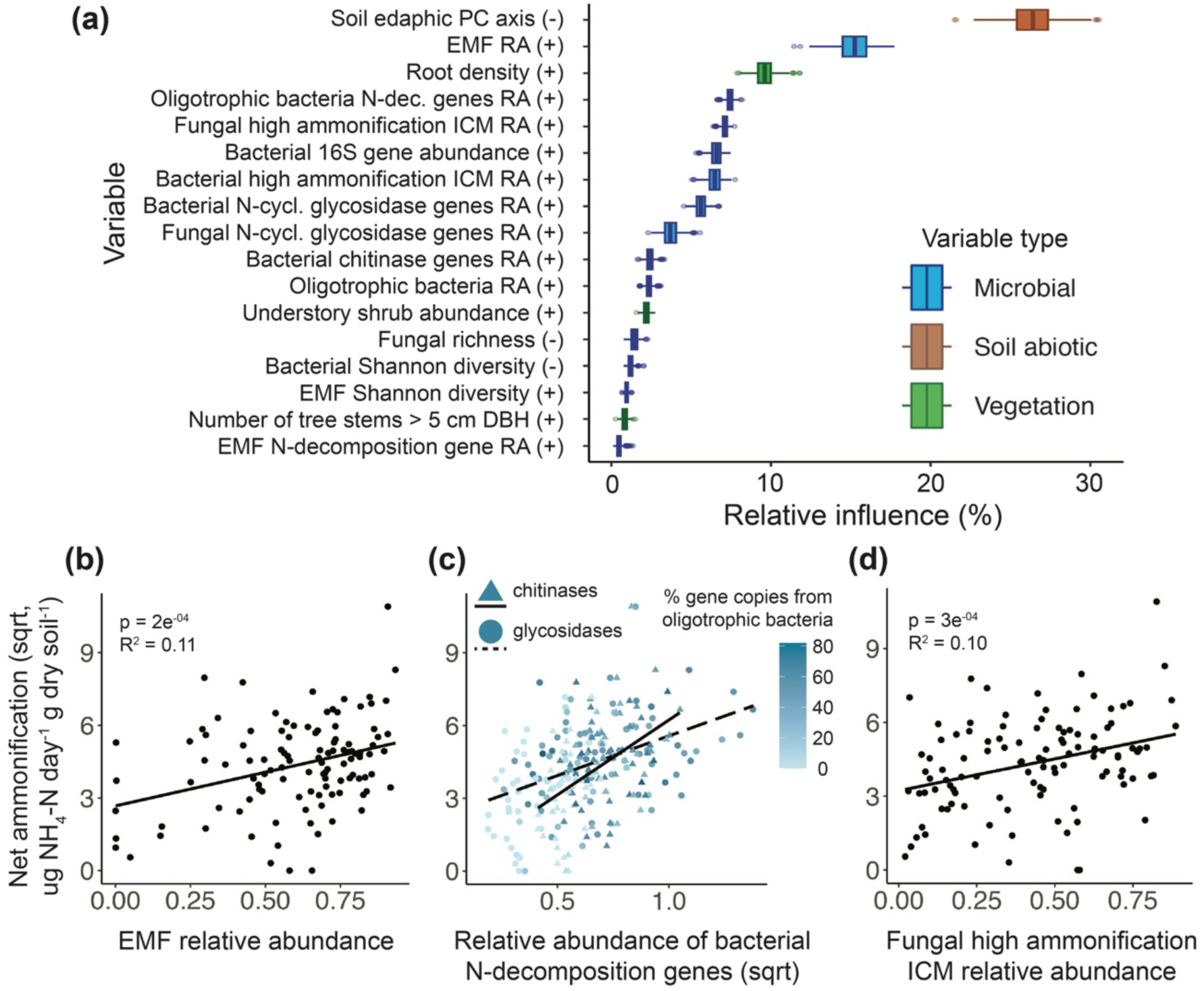
**(a)** Box plots show the relative contributions of each variable to explaining net ammonification rates across 1000 replicate model runs of Boosted Regression Tree modeling. The direction of association with net ammonification is shown in parentheses after the variable name. Linear mixed effects models show that net ammonification is correlated positively with the relative abundances (RA) of **(b)** ectomycorrhizal fungi (EMF), **(c)** all bacterial chitinase (p = 2e^-08^, R^2^ = 0.24) and N-cycling glycosidase genes (hydrolysing O- and S-glycosyl compounds, p = 6e^-08^, R^2^ = 0.23), and oligotrophic bacterial chitinase (p = 2e^-05^, R^2^ = 0.13) and N-cycling glycosidase genes (p = 1e^-04^, R^2^ = 0.11), and **(d)** the fungal high ammonification indicator co- occurrence module (ICM).

The most important microbial variables contributing to the BRTs explaining net nitrification were all related to bacteria: the relative abundances of the bacterial high nitrification ICM (Fig. 2a,b), nitrifying bacteria (Fig. 2a,c), and copiotrophic bacteria (Fig. 2a,d). In contrast to our prediction, nitrifying genes from nitrifying bacteria were not found in our dataset. Instead, nitrification rates correlated most strongly and negatively with the total relative abundance of bacterial nitrate reductase, the main enzyme involved in denitrification (Fig. 2a, Appendix S1: Fig. S9). Nitrification was also negatively associated with factors related to EMF, specifically EMF Shannon diversity and the relative abundance of medium-distance exploration EMF (Fig 2a, Appendix S1: Fig. S9). For both net ammonification and net nitrification, the highest-ranked microbial variables in the BRTs did not necessarily have the strongest linear correlations with the process rate (Fig.s 1 and 2, Appendix S1: Fig.s S8 and S9), indicating that these variables best explain net nitrification through non-linear relationships.

**Fig. 2.**
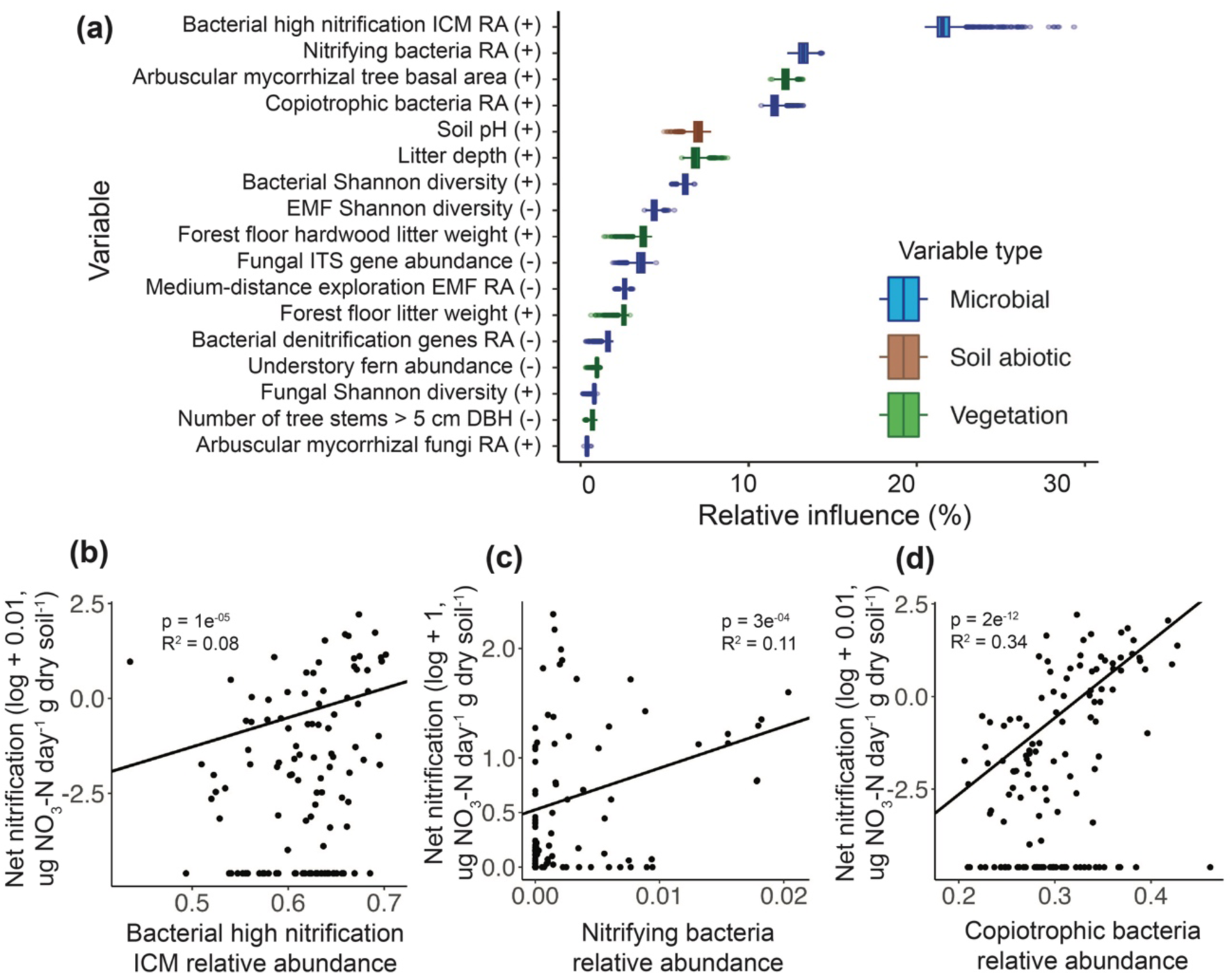
**(a)** Box plots show the relative contributions of each variable to explaining net nitrification rates across 1000 replicate model runs of Boosted Regression Tree modeling. The direction of association with net nitrification is shown in parentheses after the variable name. Zero-inflated linear mixed effects models show that net nitrification is correlated positively with the relative abundances (RA) of **(b)** the bacterial high nitrification indicator co-occurrence module (ICM), **(c)** nitrifying bacteria, and **(d)** copiotrophic bacteria. Other abbreviations are ectomycorrhizal fungi (EMF) and diameter at breast height (DBH).

The most important variable contributing to the models explaining net phosphate change was not EMF P-decomposition genes as predicted, but the relative abundance of all fungal P-cycling oxidoreductase genes. These genes had the highest contribution to BRTs and had a positive relationship with rates of net phosphate change (Fig. 3a,b). Few factors overall correlated with net phosphate change (Appendix S1: Fig. S10), but most were relative abundances of functional genes: in addition to fungal P-cycling oxidoreductase genes, the relative abundance of hydrolase genes (acting on P-bonds) from oligotrophic bacteria and the relative abundance of bacterial hydrolase genes (acting on C-P bonds) positively correlated with net phosphate change (Fig. 3a, Appendix S1: Fig. S10).

**Fig. 3.**
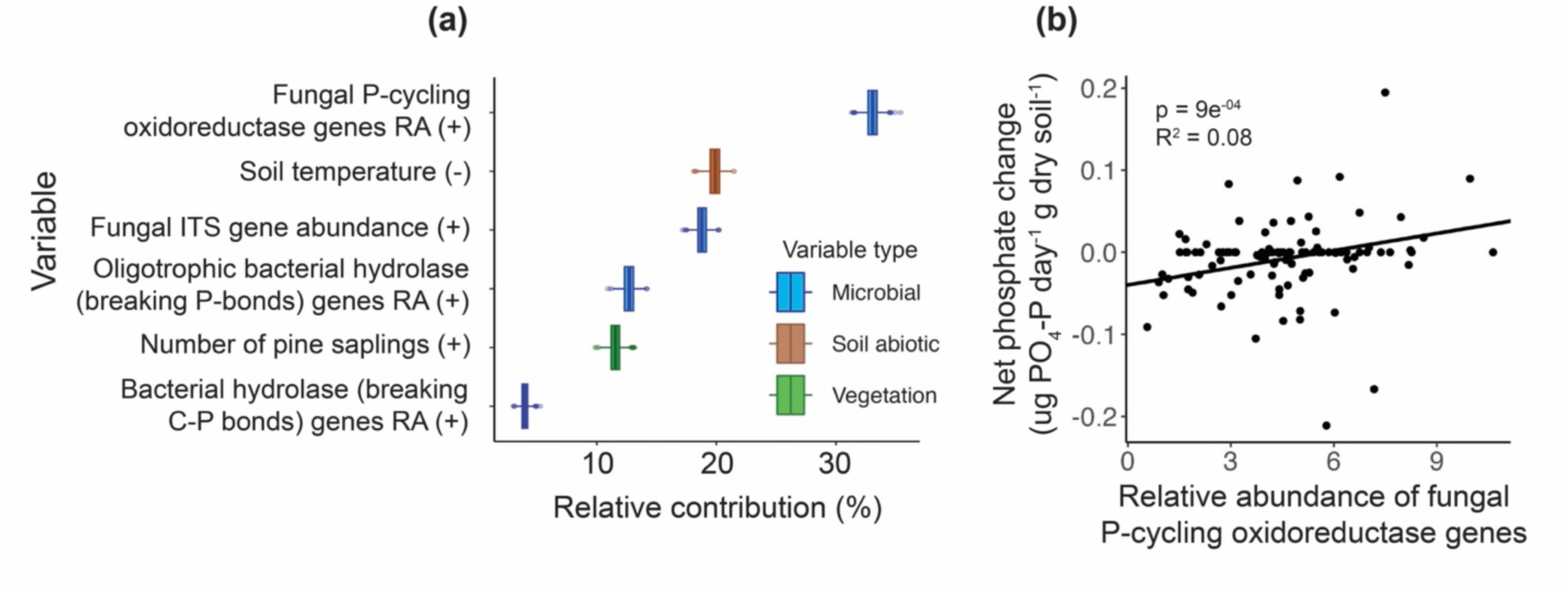
**(a)** Box plots show the relative contributions of each variable to explaining net phosphate change rates across 1000 replicate model runs of Boosted Regression Tree modeling. The direction of association with net phosphate change is shown in parentheses after the variable name. **(b)** Linear mixed effects model shows that net phosphate change is correlated positively with the relative abundance (RA) of fungal P-cycling oxidoreductase genes.

In contrast to our second hypothesis, microbial metrics did not explain more variation in net ammonification rates than abiotic factors, but they were still included in the best multivariate linear models that explained the most variation in net ammonification (Table 2). These models included a soil edaphic variable (PC axis) and either the relative abundance of EMF or the fungal high ammonification ICM (Table 2). When removing microbial variables from their respective models, the total variance explained by the model did not decrease, but the model fit declined (AIC change of +64). This finding is corroborated by the BRTs, where the soil edaphic PC axis was by far the most important variable explaining variation in net ammonification rates (Fig. 1a).

**Table 2.**
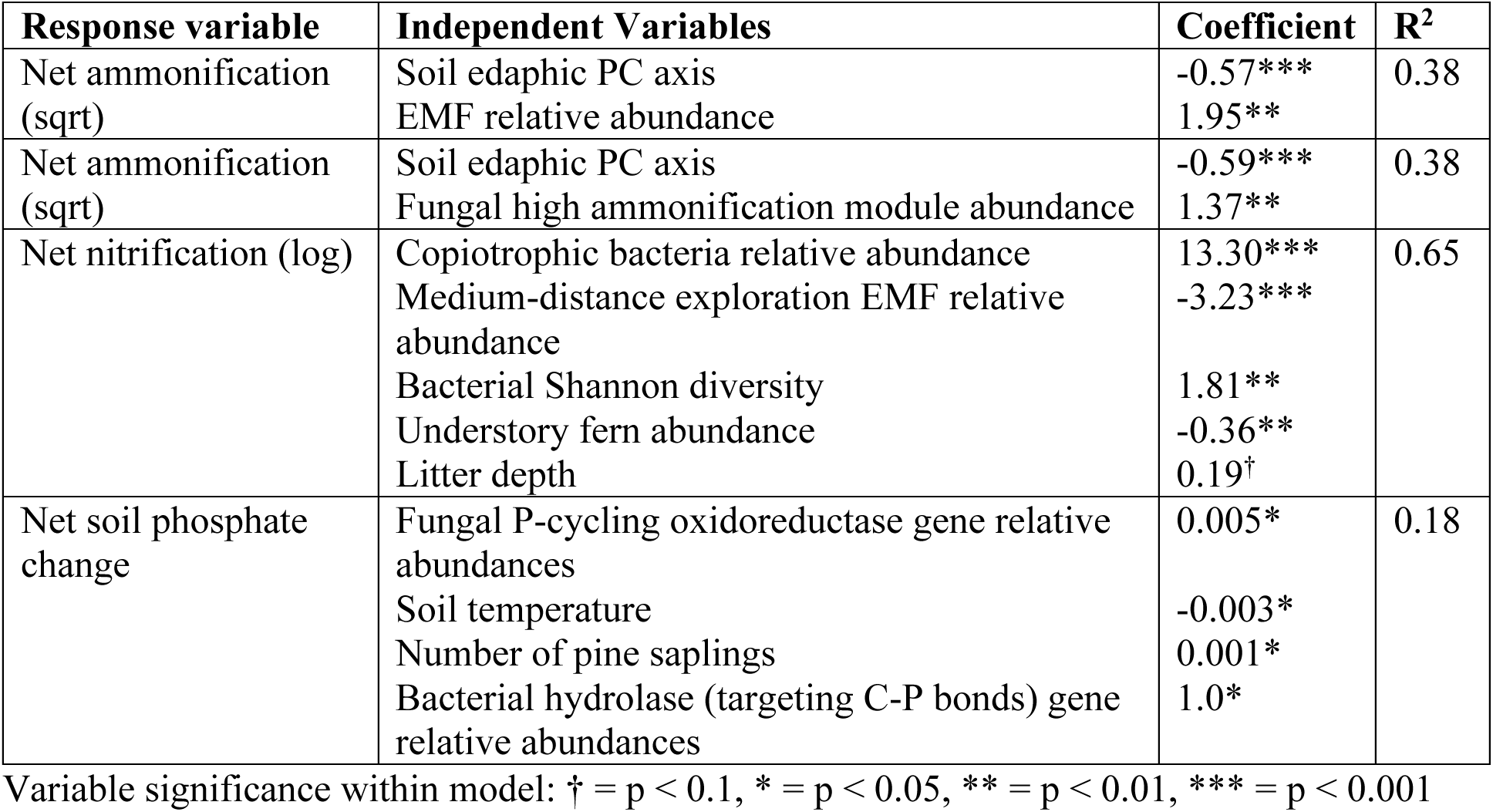
Best-fit multivariate linear models explaining soil nutrient cycling rates based on minimum AIC value.

By contrast, microbial variables explained more variation than plant and soil edaphic factors in models of net nitrification and phosphate change rates. The best model for net nitrification included only microbial and plant variables, specifically the relative abundances of copiotrophic bacteria and medium-distance exploration EMF, bacterial Shannon diversity, fern abundance, and litter depth as explanatory variables (Table 2). Microbial variables explained the most variation in the linear model for net nitrification: when microbial variables are removed from the model, the R^2^ decreased by 0.46, or 71% of the total variance explained by the best model (Table 2). Vegetation factors were also important in both the BRTs and the multivariate linear model, especially AM tree basal area (Fig. 2a) and litter depth (Fig. 2a, Table 2).

Net phosphate change was best explained by a combination of microbial, plant, and soil edaphic factors: the best multivariate linear model included the relative abundances of fungal P-cycling oxidoreductase genes and bacterial hydrolase (targeting C-P bonds) genes, soil temperature, and the number of pine saplings (Table 2). When microbial variables are removed from the model, the R^2^ decreased by 0.11, or 61% of the total variance explained by the best model, yet this best model explained relatively low variance in net phosphate change (18%, Table 2). The results from the BRTs similarly showed that the relative abundance of fungal P-cycling oxidoreductase genes was more important than soil and vegetation variables, but soil temperature is still important as the second-highest contributing variable (Fig. 3a). Additionally, litterfall chemistry variables (magnesium and molybdenum concentrations) were important in the transect-level models explaining net phosphate change (Appendix S1: Table S2).

## DISCUSSION

Despite decades of soil microbial ecology research, the aspects of microbial communities that are most useful for explaining soil nutrient cycling have remained elusive. Our study addresses this outstanding ecological question, finding that functional guild relative abundances and indicator co-occurrence modules were the most important microbial variables explaining N cycling, while whole-fungal community P-cycling gene abundances were the most important microbial variable explaining soil P cycling. Net ammonification was best explained by soil edaphic factors, but microbial variables explained more variance in net nitrification and phosphate change than plant and soil abiotic variables, suggesting that microbial activity is a key regulator of these processes. These results suggest a new framework for the roles of different microbial, plant, and soil abiotic factors in specific N- and P-cycling processes (Fig. 4).

**Fig. 4.**
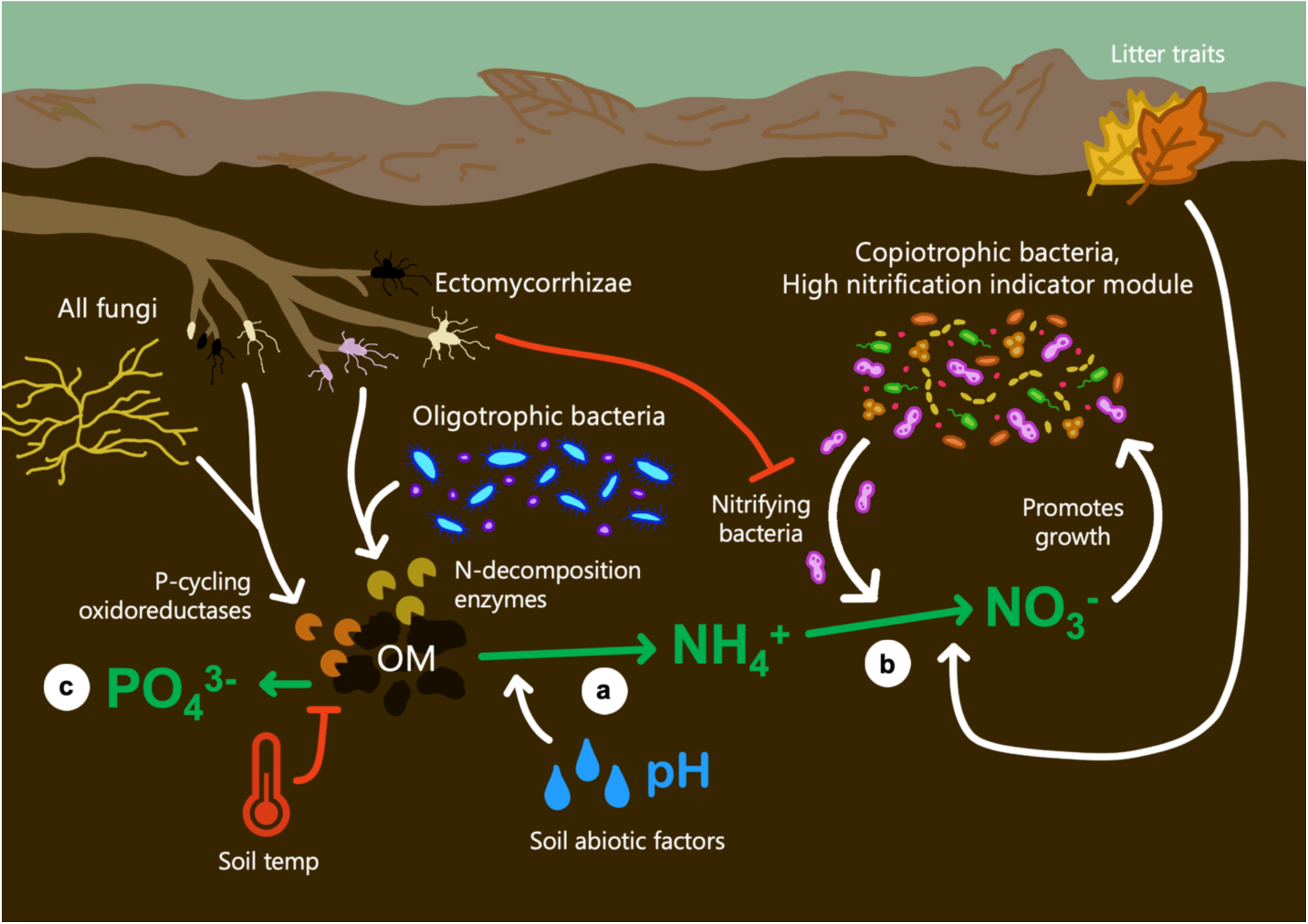
Hypothetical drivers and controls over soil nutrient cycling based on study results. Results from our field system analysis suggest that: **(a)** Net ammonification is driven by N- decomposition enzymes produced by ectomycorrhizal fungi and oligotrophic bacteria, which is regulated by edaphic factors like soil organic matter (OM) content, moisture, and pH. **(b)** Nitrification is driven by nitrifying bacteria within the guild of copiotrophic bacteria and the high nitrification indicator co-occurrence module, which in turn promotes the growth of other bacteria within these groups. Nitrification is also influenced positively by litter traits like litter depth, and negatively by ectomycorrhizal fungi, which inhibit the growth and activity of nitrifying bacteria, potentially through competition for ammonium, decreasing net nitrification rates. **(c)** Net phosphate change is driven by P-cycling oxidoreductases produced by a wide range of soil fungi, and is inhibited by soil temperature.

### Microbial metrics explaining nutrient cycling rates

The microbial metrics explaining the most variation in N-cycling processes were consistently related to specific microbial functional groups: EMF and oligotrophic bacteria for net ammonification, and nitrifying and copiotrophic bacteria for net nitrification. The relative abundances of EMF, oligotrophic bacteria, and their N-decomposition genes were all positively associated with net ammonification (Fig. 1a-c, Appendix S1: Fig. S8), indicating that EMF and oligotrophic bacteria may drive ammonification by producing N-decomposing enzymes (Fig. 4a). EMF and oligotrophic bacteria are characteristic of slow nutrient-cycling systems, where most nutrients are found in organic forms (Fierer et al. 2007, Phillips et al. 2013, Ho et al. 2017), such as in the nutrient-limited northern temperate forests represented by our study sites (Groffman et al. 2018). EMF may break down organic matter in search of N rather than C (Talbot et al. 2015, Nicolás et al. 2019) and can produce high levels of organic matter- and N- decomposing enzymes (Talbot et al. 2008, Bödeker et al. 2014, Lindahl and Tunlid 2015, Zak et al. 2019) such as CAZymes, chitinases, ureases, and peroxidases (Argiroff et al. 2022, Maillard et al. 2023, Zhang et al. 2023, Auer and Buée et al. 2024). While little work has examined the N- decomposition potential of oligotrophic bacteria as a group, there is evidence that marine oligotrophic bacteria target N-rich molecules through leucine amino peptidase activity (Rath et al. 1993). In addition, the oligotrophic bacterial phyla abundant in our study sites, such as *Acidobacteria*, have high genomic potential to degrade complex N-containing polysaccharides (Belova et al. 2018, Zhou et al. 2019, Kalam et al. 2020). Our findings add to this body of evidence that EMF and oligotrophic bacteria may drive the decomposition of N in soils of these temperate forest ecosystems, and, to our knowledge, is the first evidence showing that oligotrophic bacterial and EMF N-cycling genomic potential correlate with N-cycling rates in ecosystems.

Net nitrification rates were best explained by taxa within the bacterial high nitrification ICM, nitrifying bacteria, and copiotrophic bacteria (Fig. 2, Fig. 4b), indicating that these taxa may be the primary drivers of nitrification processes in these temperate forest soils. Our observations are consistent with the findings that nitrification is largely performed by bacteria and archaea (Stein and Klotz 2016). Known nitrifying bacteria are included within the copiotrophic bacteria group and high nitrification ICM (Appendix S1: Fig. S3), but there may be more taxa in these groups that perform nitrification than we were able to annotate based on literature and genomic databases: there were multiple samples with positive net nitrification rates that contained no nitrifying bacteria based on our annotations. This may have occurred because many bacterial ASVs (47%) were not assigned taxonomy at the genus level, which is needed to identify many taxa as nitrifiers, some bacterial lineages may contain unidentified nitrifiers (Liao et al. 2024), or nitrification in these samples may have been performed by archaea or other microbes (Nicol et al. 2008, Isobe et al. 2015). It may also be the case that copiotrophic bacteria and taxa in the high nitrification ICM grow and thrive in the environmental conditions created by nitrifiers (i.e., high inorganic nitrogen availability). The high nitrification ICM may be especially important because it is calculated via correlation networks between taxa (Langfelder and Horvath 2008), capturing associations between nitrifying bacteria and taxa that may depend on nitrifying bacterial activity (McClure et al. 2022). Net nitrification was also negatively associated with variables related to EMF, especially medium-distance exploration EMF, potentially due to competition for inorganic N between mycorrhizae and nitrifying bacteria and archaea (Fig. 4b, Tatsumi et al. 2020, Sun et al. 2023). Medium-distance exploration EMF have longer hyphal lengths and increased nutrient foraging distance compared to contact or short-distance exploration EMF types (Agerer 2001, Chen et al. 2018) and are hypothesized to have limited capacity to degrade organic N (Hobbie and Agerer 2009), so they may take up large quantities of ammonium to compete with nitrifiers.

In contrast to net ammonification and nitrification, net phosphate change was not linked to any specific groups of microbial taxa, but was instead linked to fungal and bacterial P-cycling oxidoreductase and hydrolase genes (Fig. 3). While the role of microbial oxidoreductases in soil P cycling has been understudied, redox reactions are being recognized as increasingly important in the soil P cycle (White and Metcalf 2007, Ewens et al. 2021, Kehler et al. 2021). These results indicate that the cumulative sum of the entire microbial (especially fungal) community’s P-targeting enzyme activity may be the primary driver of P release from soil organic matter (Fig. 4c).

### Relative contributions of microbial, vegetation, and soil abiotic factors

Despite the widespread measurement of soil edaphic variables to explain nutrient cycling rates in ecosystems (Elrys et al. 2021), soil properties were only key to explaining two of the three nutrient cycling rates we measured. For net ammonification, the soil edaphic PC axis (that includes pH, % SOM, % soil moisture, soil temperature, and organic layer depth) explained by far the most variance in net ammonification of any variable (Fig. 1a, Table 2), indicating that these factors together are the primary controls over release of ammonium from SOM (Fig. 4a). In contrast, for net phosphate change, soil temperature was the only soil abiotic variable included in the BRTs or multivariate linear model (Fig. 3a, Table 2). Soil P availability globally is negatively associated with temperature, potentially due to increasing plant P uptake or reducing microbial P-decomposition activity (Unger et al. 2010, Hou et al. 2018, Hu et al. 2022), indicating that temperature is a strong control over the release of phosphate in soils (Fig. 4c). Some microbes exhibit a tradeoff between stress tolerance (for example, heat tolerance) and nutrient acquisition activity (Malik et al. 2020), which could result in decreased P cycling activity under stressful environmental conditions. Surprisingly, for net nitrification, no soil abiotic variables were included in the multivariate linear model or were among the top variables contributing to the BRT (Figure 2a, Table 2). pH was the only soil abiotic variable that significantly correlated with net nitrification (Appendix S1: Fig. S9), which may impact nitrification rates through affecting the abundances of pH-sensitive nitrifying microbes (Stempfhuber et al. 2015, Nicol et al. 2008). Because nitrification is a primarily microbe-mediated process (Stein and Klotz 2016), microbial variables may overshadow soil variables that affect nitrification indirectly.

Several different plant factors explained variance in nutrient cycling processes, some of which were unexpected. In line with previous findings, the basal area of trees that associate with arbuscular mycorrhizal (AM) fungi (Fig. 2a) and litter depth (Fig. 2a, Table 2) were especially important in explaining net nitrification, and may affect nitrification rates indirectly through litter quality (Fig. 4b). AM trees often have higher litter quality (e.g., low litter C:N, Laughlin 2011) and increase soil inorganic N concentrations (Phillips et al. 2013, Lin et al. 2017, Lin et al. 2018), as well as nitrifier microbe abundances (Teutscherova et al. 2019). For net ammonification, root density was the third-most influential variable in the BRTs (Fig. 1a), which may impact net ammonification by controlling the abundance and composition of EMF (Peay et al. 2011), or by increasing the availability of C-rich root exudates, priming microbial decomposer activity (Dijkstra et al. 2013, Yin et al. 2018). Surprisingly, for net phosphate change, the number of pine saplings was included in both the BRTs and the final linear model as an important explanatory variable (Fig. 3a, Table 2). Because pine saplings contributed negligible pine litter to the forest floor (Appendix S1: Fig. S2), but can bring a unique EMF community into forests (Policelli et al. 2020), these results indicate that P cycling may be shaped by the EMF community. Early-successional pine-specific fungi (e.g., EMF communities unique to non-mature pines) may promote P-decomposition (Delucia et al. 1997), potentially through the direct production of P-cycling enzymes and low-molecular-weight organic anions (Courty et al. 2006, Alvarez et al. 2012, Wang and Lambers 2020, Wang et al. 2021) or by hosting bacterial communities with high phosphatase activity (Yuan et al. 2024).

Few factors overall explained net phosphate change and even the best multivariate linear model explained a low percent of variance (18%, Table 2). This may be because of low soil inorganic P levels at our sites, where most of our samples had zero net phosphate change and low overall variance in rates. Northern temperate forests often have low concentrations of soil inorganic P, such that all P liberated from organic matter may be taken up by microbes and plants nearly immediately (Sohrt et al. 2017, Forstner et al. 2019), making it difficult to quantify gross rates of phosphate release from organic matter. However, P-cycling in temperate forests is understudied and net rates of phosphate change are almost never quantified. More studies examining the biotic drivers of P-cycling rates, especially gross rates of inorganic P release or uptake, will improve our understanding of the mechanisms behind P cycling in temperate forests.

Our work showed that, despite their caveats, commonly measured microbial variables from amplicon sequencing are linked to soil nutrient cycling rates in northern temperate forests. Instead of measuring single marker genes (via qPCR), estimating groups of genes – as we did in this study – may better capture potential microbial metabolic pathways involved in ecosystem processes. However, many microbial nutrient cycling pathways remain unannotated (Albright et al. 2019) and it is unclear whether functional gene abundances always reflect gene expression and enzyme activity. PICRUSt2 or similar methods for fungi (Anthony et al. 2022) likely under- estimate certain gene groups, including key nitrification genes that were not found in our dataset, which has also been observed in other studies (Tatsumi et al. 2023). For fungi, the relatively small number of available fungal genomes means we must assign most gene counts at the genus level, which ignores species-level variation (Bödeker et al. 2014). Because of these issues, using gene expression data, such as meta-transcriptomic data, for key functional groups may be better correlated with process rates than genomic potentials. On the other hand, gene counts can reflect accumulated functional potential over ecological and evolutionary timescales and our analyses show that even conservative estimates of functional gene abundances can explain significant variation in nutrient cycling rates. Estimated gene counts may be good metrics to predict flux rates in areas where they haven’t been measured yet, but soil microbial community composition data is available.

### Conclusions

Through the first comprehensive test comparing a broad range of microbial community traits, we found that the relative abundances of functional guilds and indicator co-occurrence modules explain soil N-cycling rates, while the relative abundance of fungal P-cycling genes explain soil P-cycling rates. These results suggest a new conceptual model for these biogeochemical processes (Fig. 4), where net ammonification is driven by soil abiotic factors and N-decomposition enzymes produced by EMF and oligotrophic bacteria, net nitrification is tightly coupled to copiotrophic bacteria and the bacterial high nitrification ICM and regulated by litter traits, and net phosphate release is driven by oxidoreductases produced by the whole fungal community and soil temperature. This analysis is a key step towards distilling the complexity of microbial communities into quantifiable terms explaining and predicting ecosystem function that could improve predictions of nutrient movement through ecosystems.

## ACKNOWLEDGEMENTS

We would like to thank members of the Bhatnagar lab for their assistance with field work, manuscript edits, and laboratory analysis, especially Chikae Tatsumi, Michael Silverstein, Zoey Werbin, Kathryn Atherton, Victoria Moscato, Katherine Sotiropoulou, Long Hong, Li Lin, and Anthony Avila. We also thank those who gave us permission to sample at the field sites, including the Harvard Forest, the Newton Massachusetts Conservation Commission, the Burlington Massachusetts select board, the Lexington Massachusetts Conservation Commission, and the Maronite Monks of Adoration in Petersham, Massachusetts. This research was funded by the Department of Energy Biological and Environmental Research awards DE-SC0020403 and DE-SC0012704 to Jennifer Bhatnagar, an NSF Graduate Research Fellowship to Corinne Vietorisz, and Research Awards from Boston University’s Graduate Student Organization and Biogeoscience program to Corinne Vietorisz.

## AUTHOR CONTRIBUTIONS

C.V., N.P., and J.M.B. established sampling sites and design. C.V., N.P., L.A., and J.M.B. conducted field sampling. C.V., N.P., A.L, and L.A. performed laboratory analysis on samples.

C.V. and J.M.B. conducted data analysis. C.V. and J.M.B wrote the first draft of the manuscript, and all authors edited the manuscript.

