## Appendix S1 for "Microbial functional guilds and genes are key to explaining soil nutrient cycling alongside soil and plant variables"

Supplemental Information

**Microbial functional guilds and genes are key to explaining soil nutrient cycling alongside soil and plant variables**

Corinne Vietorisz, Nahuel Policelli, Abigail Li, Lindsey Adams, Jennifer M. Bhatnagar

SECTION S1: METHODS

***Site descriptions***

We selected six field sites: three in suburban forests in Eastern Massachusetts, including Landlocked Forest (Lexington, MA, 42.484, -71.229), Whipple Hill Conservation area (Burlington, MA, 42.437, -71.185), and Hammond Woods (Newton, MA, 42.327, -71.175), and three in the heavily forested areas of central Massachusetts in or adjacent to the Harvard Forest Long Term Ecological Research site, two within Petersham, MA (42.461, -72.166 and 42.466, -72.213) and one within Phillipston, MA (42.551, -72.174). Each site contains 4 forest stand types, all within 2.5 km of each other. Pine forests were dominated by mature white pines along the transect, with minimal pine saplings in the understory. Mature hardwood forests were dominated by mature hardwood trees, primarily *Quercus,* along the transect, with no pines present. Encroachment sites contained no mature pines, but had encroaching white pine saplings (< 5 cm Diameter at Breast Height, DBH) in the understory along the transect. Mixed forests contained a mix of mature white pine trees and mature hardwood trees, with minimal pine saplings in the understory. *Pinus strobus* was the only pine species present at all sites, while the dominant hardwood species present across all sites were *Quercus rubra* (51% of all hardwood trees across all sites)*, Acer rubrum* (16%)*, Acer saccharum* (9%) and *Betula lenta* (5%). These forest types were not used for differential analysis of response variables between forest types, but provided variation in soil microbial community and vegetation characteristics.

***Soil variables***

During soil collection, at each sampling point, the litter layer was removed before sampling and a 10 cm x 10 cm soil brownie was collected to a depth of 6 cm by cutting into the soil with a knife. To calculate soil net ammonification and nitrification, bulk soil samples were collected in mid-July at every sampling point (204 total), and 144 bulk soil samples were buried at the 0 m, 15 m, and 60 m A and B sampling points. Four buried bags were unable to be recovered at the end of the 4-week incubation, so 140 buried soil bags were collected. Within 48 h of sample collection, soil was sieved through a 2 mm sieve and 10 g of soil from each sample was used for inorganic nitrogen extraction with 2M potassium chloride (Caron et al. 2023). The extracts were frozen at -20 °C for subsequent colorimetric analysis. Soil extracts were analyzed colorimetrically for $NH4^+$ and $NO3^-$ concentrations using a colorimetric method (Caron et al. 2023). Net rates of ammonification and nitrification per gram dry soil per day were calculated as the differences in ammonium and nitrate concentrations between the initial soil sample and the buried bag soil sample divided by the total days incubated.

To calculate net soil phosphate change in July – August 2022, we buried 144 soil samples in the same locations as the 2021 buried bags. 8 bags were destroyed or unable to be recovered at the end of the 4-week incubation, so 136 buried soil bags were collected. Within 72 h of sample collection, soil was sieved through a 2 mm sieve and 1.5 g of soil was used for Olsen P extraction (Olsen et al. 1954, Frank et al. 1998). The extracts were frozen at -20 °C for subsequent analysis. Soil extracts were analyzed colorimetrically for $PO_4^{3-}$ using a novel microplate method. Soil extracts were pipetted into a 96-well microplate, including phosphate standard solutions ranging from 0 – 10 ppm $PO_4$-P (Ricca Chemical). An acid molybdate solution containing ammonium molybdate, antimony potassium tartrate, sulfuric acid, and ascorbic acid was added to the extracts and standards, incubated at room temperature for 10 minutes, and the resulting color intensity was quantified in a microplate reader (Synergy H1, Biotek) at an absorbance wavelength of 882 nm (Frank et al. 1998). Net rates of phosphate change per gram dry soil per day were calculated as the differences in phosphate concentration between the initial soil sample and the buried bag soil sample divided by the total days incubated.

***Vegetation variables***

To assess leaf litter composition, a 20 cm x 20 cm square was placed upon the forest floor in between the A and B sampling points, a knife was used to cut along the edges of the square, and all litter within the square was collected. Within 12 h of collection, the litter was dried at 65 °C for 48 h and then weighed for total litter mass. Forest floor litter samples were then sorted by hand into hardwood litter, pine litter, and other litter. The weight of litter in each category was recorded for each sample.

To measure root density, one 10.2 cm deep, 5.1 cm diameter soil core was taken at each distance between the A and B sampling points using a slide hammer core. All roots that could not pass through a 2mm sieve were extracted from the core, dried for 48 h at 60°C and weighed. Root density was calculated as the total dry weight of roots per cubic cm of soil.

To assess litterfall characteristics, falling leaf litter from the forest interior was collected once per transect (24 total transects) for one full year from mid-September 2021 until mid-September 2022 via litter collection baskets. Litter collection baskets were placed on the forest floor. Each litter collection basket was 28cm x 33cm x 33cm and lined by 1.5 mm mesh to keep the leaf litter from touching the forest floor and prevent the collection of water. Two leaf litter collection baskets were installed at the A and B sampling points at either 30 m or 60 m from the forest edge at each transect, depending on which distance was most representative of the whole-transect vegetation. Litterfall was collected at multiple time points at all transects to capture both hardwood and pine litterfall timing: late October 2021, mid-November 2021, early December 2021, late May 2022, early August 2022, and mid-October 2022. During each collection, the leaf litter from the two baskets at each transect was pooled, dried for 48 h at 65°C, and weighed. To obtain litterfall chemistry, leaf litter from the peak litterfall collection (mid-November 2021) was homogenized and ground into a fine powder for total C and N analysis according to the same protocols as soil total C and N. Litterfall total micronutrient concentrations (in micrograms per gram plant material) was measured on air-dried and homogenized litterfall at the Ohio State Service Testing and Research Laboratory via the EPA 3051A acid digestion (U.S. EPA 2007).

To assess tree community composition, the diameter at 1.4 m from the ground (Diameter at Breast Height, DBH) of all trees was measured for every tree > 5cm DBH. Each tree was identified to genus and to species if possible. In each plot, the number of white pine saplings under 5 cm DBH was counted to assess the density of white pine encroachment. To survey understory vegetation composition at each sampling point, we established 1m x 1m quadrats around each A and B sampling point. In each quadrat, the understory cover was grouped into categories based on the vegetation growth forms at our sites: broad-leaf herbs, grassy herbs, ferns, shrubs, vines, moss, white pine seedlings, oak seedlings, other tree seedlings, exposed leaf litter, bare soil, and rock. The coverage of each category within the plot was assessed using the Braun-Blanquet method (Braun-Blanquet 1932, Matteucci and Colma 1982).

***Microbial DNA amplicon sequencing and bioinformatics***

After DNA amplification, amplicon quality was checked via agarose gel electrophoresis, then amplicons were cleaned with the Just-a-Plate 96 PCR Purification and Normalization Kit (Charm Biotech, MO) and quantified using the Qubit HS-dsDNA kit (Invitrogen, Carlsbad, CA). To prepare libraries for sequencing, 16S and ITS amplicons were each pooled at 25 ng of DNA per sample, then both 16S and ITS amplicons were combined into a single library for sequencing, with two libraries in total: one library for all “A” sampling points, and one library for all “B” sampling points. Each library was subject to 250 base pair (bp) paired-end sequencing on an Illumina MiSeq run at the Tufts Genome Sequencing Core facility.

While running the dada2 pipeline to process DNA sequence data, for ITS sequences, primers were trimmed with the R package cutadapt (Martin et al. 2011). Forward ITS reads were trimmed at 250 bp, reverse reads were trimmed at 210 bp, and the minimum read length was set to 50 bp. For 16S sequences, the minimum read length was set to 100 bp, and no base pairs were trimmed aside from trimming the primers because quality scores largely remained above 30 for the entire sequence. Taxonomy was assigned to ASVs using the naive Bayesian classifier method (Wang et al. 2007) in combination with the UNITE database (v. 9.0) as the reference for fungal ITS ASVs (Abarenkov et al. 2024, Nilsson et al. 2019), and the SILVA database (release 138.1, Quast et al. 2013) as the reference for bacterial ASVs. Across 194 soil samples, 17,021,506 bacterial sequences were detected, of which 10,465,199 remained after all filtering steps (52,082 +/- 34,758 sequences per sample), containing 20,204 unique ASVs. 5,044,748 fungal sequences were detected, of which 2,859,290 remained after filtering (14,514 +/- 5,669 sequences per sample), containing 9,309 unique ASVs. For ITS sequences, 13 low-read samples (with less than 8000 reads after all filtering steps) were removed from analysis, resulting in a total of 181 samples with ITS sequence data. For 16S sequences, 3 samples were determined as outliers due to sequencing errors or contamination were removed, resulting in a total of 191 samples with 16S sequence data.

***Microbial community characteristics***

For the estimated functional gene relative abundances, 438 bacterial ASVs out of 20,204 were assigned E.C. numbers, comprising 61% of the total 16S reads in our dataset. 2,494 fungal ASVs of the 9,309 in our dataset were assigned genome annotations at either the genus or species level, comprising 55% of the total ITS reads in our dataset. 66 fungal ASVs were assigned a species-level genome annotation match and 2,428 fungal ASVs in our dataset were assigned a genus-level genome annotation average. ASVs without genus-level taxonomic assignments were not included in the analysis.

To calculate the relative abundances of indicator co-occurrence modules, we ran a Weighted Gene Correlation Network Analysis (WGCNA) using the 'WGCNA' R package (Langfelder and Horvath 2008). We input net ammonification, net nitrification, and net phosphate change rates as the trait data. All ASVs that occurred in less than 2 samples were removed, leaving 2,688 fungal ASVs and 6,459 bacterial ASVs remaining. The R package 'flashClust' (Langfelder and Horvath 2012) was used for detecting outliers of samples with the clustering method "average." No outliers were detected for ITS or 16S. Modules of frequently co-occurring ASVs were identified using the dynamic tree cut method based on dissTOM hierarchical clustering, with deepSplit = 2 and minModuleSize = 4. Modules with dissimilarity coefficients of 0.9 were merged for ITS and coefficients of 0.8 were merged for 16S, resulting in 76 total modules for ITS and 48 total modules for 16S. The moduleTraitCor() function was used to correlate rates of nutrient cycling with co-occurrence module principal components. Significant correlations between a co-occurrence module first principal component and net ammonification, net nitrification, or net phosphate change were identified with the function corPvalueStudent(). Modules with highly significant relationships with nutrient cycling rates ($p =< 0.001$) were selected. For bacteria, 7 modules were highly significant with net ammonification, 13 with net nitrification, and 4 with net phosphate change. For fungi, 4 modules were highly significant with net ammonification, 10 with net nitrification, and 7 with net phosphate change. A list of ASVs that most strongly belonged in each module ($kME > 0.66$) was compiled. Then, for all the modules that were highly significantly positively or negatively associated with each nutrient cycling metric, the cumulative relative abundances were summed of all ASVs with a kME > 0.66 in those modules (indicating that a taxa is central to the network, Fig.s S3 and S4), using ASV relative abundances. This created a new metric of "indicator co-occurrence module abundance" for bacteria and fungi for high ammonification, low ammonification, high nitrification, low nitrification, high phosphate change, and low phosphate change modules.

***Statistical methods***

Explanatory variable selection

To reduce the number of input explanatory variables in model selection analyses, we selected the following variables. For microbial functional guilds (and relative gene abundances from functional guilds) that were inversely correlated because of the compositional nature of microbial amplicon sequencing data (eg, EMF and saprotrophic fungi) and functional guilds where one is a subset of the other (eg, EMF and medium-distance exploration EMF), we input whichever variable had the strongest correlation with the process rate. N-decomposition gene relative abundances were all highly correlated with each other, so we input the two N-decomposition gene groups from both bacteria and fungi that had the strongest correlations with net ammonification. N-decomposition gene abundances from functional guilds (i.e., from oligotrophic bacteria and EMF) were all highly correlated with each other and had similar correlation strengths with net ammonification, so we created two new variables summing all N-decomposition gene abundances from oligotrophic bacteria and from EMF, separately. For indicator co-occurrence modules, either the "high" or "low" module was picked for both fungi and bacteria for each process (whichever one had the stronger correlation). Additionally, because many soil abiotic variables co-correlated with each other, a principal component analysis (PCA) was run on the soil abiotic variables % SOM, soil pH, soil temperature, soil % moisture, and organic layer depth (Fig. S5), and a new variable "edaphic PC1" was created by extracting the first principal component. Soil total C and total N were not included in the edaphic PCA because of especially strong correlations with % SOM and each other, so % SOM was selected to represent soil nutrient properties. Most soil edaphic variables were correlated with net ammonification, and were correlated with each other, so we input the edaphic PC1 into the ammonification model selection as the only soil abiotic variable.

Because the zero-inflated linear model selection for net nitrification is intensive to complete manually, we reduced the number of input variables relative to those input into the BRT by removing highly co-correlated variables. We ran a PCA on the set of BRT input variables (Fig. S7), and when there were clear groups of variables that strongly correlated with each other in PCA space, only the variable with the strongest correlation with net nitrification (Fig. S9) was selected. If two variables had similar correlation strengths, then the one with the most biological relevance was selected. The only exception was for the relative abundance of nitrifying bacteria and soil pH, which were both kept in the model selection process because of their especially strong correlations with net nitrification (Fig. S9). This yielded 11 input variables into the model selection process (Fig. S7).

For the transect-level multivariate linear models, explanatory variable selection was performed as described in main text and the first paragraph of this section. The only exceptions were to reduce the number of input variables for net ammonification and net nitrification: for net ammonification, all fungal and bacterial N-decomposition gene relative abundances had similar correlations with net ammonification, so one fungal and one bacterial N-decomposition gene variable was created by summing all bacterial or fungal N-decomposition relative gene abundances. For net nitrification, multiple litterfall micronutrient concentrations correlated significantly and positively with net nitrification, so the litterfall micronutrient with the strongest correlation (litter sulfur content) was chosen as the only litterfall micronutrient variable to reduce the number of input predictors. This process resulted in 10 input variables for net ammonification, 17 for net nitrification, and 10 for net phosphate release.

Model construction and selection

The boosted regression trees (BRTs) modeling net ammonification and nitrification were fit using the gbm.step() function, which identifies the optimal number of trees for the model to build that minimizes the predictive deviance (Elith et al. 2008). We began using baseline parameters, and then adjusted the interaction depth, learning rate, resampling fraction ("bag fraction") and step size to minimize the predictive deviance following Elith et al. (2008), Allen et al. (2017), and Leathwick et al. (2006). The BRT modeling net phosphate release was fit using the gbm.fix() function with the number of trees set to the recommended minimum of 1000, and we adjusted parameters to minimize predictive deviance (Elith et al. 2008). For all three nutrient cycling processes, the BRT interaction depth was set to 2, the learning rate set to 0.001, the bag fraction set to 0.75, the step size set to 50, and cross-validation folds set to 10. Because there is slight variation in the predictor variable contributions in each run, we ran 1,000 replicate model runs for each BRT. The mean number of trees fitted across these 1,000 runs was 2,848 for net ammonification and 7,866 for net nitrification.

When running the zero-inflated linear models explaining net nitrification, the 'DHARMa' package was used to test for zero-inflation (Hartig 2022). Within the models, parameters were set: ziformula = ~ 1, family = Gamma(link = "log"). A pseudocount of 0.01 was added to net nitrification due to the log transformation, except for when the relative abundance of nitrifying bacteria was the sole explanatory variable—in this case, a pseudocount of 1 was added to avoid the production of nonexistent error values. To construct the best-fit multivariate model explaining net nitrification, we performed stepwise model selection manually on zero-inflated generalized linear mixed models using the 'glmmTMB' command (Brooks et al. 2017). A full model was constructed that included all 11 input variables, then one variable at a time was removed from the model, and the Akaike Information Criterion (AIC) of each was assessed. The variable removed from the model with the lowest AIC was eliminated, and the selection process was repeated until only one variable remained. The model with the lowest AIC from all rounds was chosen as the final best fit model.

During linear multivariate model selection for the transect-level models using Mallow's Cp score, the only exception from the model selection process described in the main text was for the transect-level nitrification model, where there were no models with a Cp score under 2 where all predictor variables were significant within the model. In this case, we chose the model with the lowest AIC and the lowest Cp score where all predictor variables were significant.

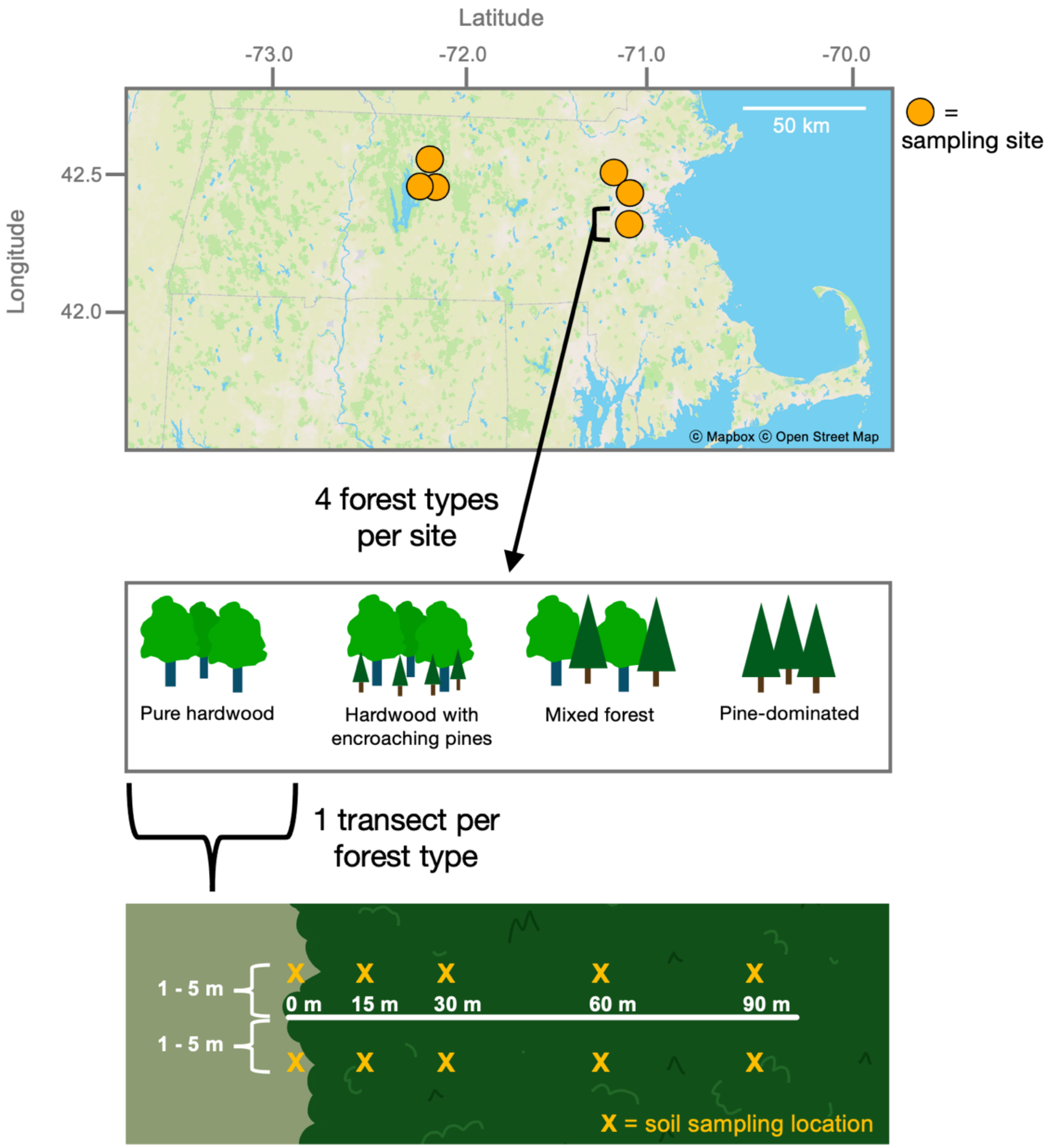


**Fig S1.** Diagram of the field sampling design.

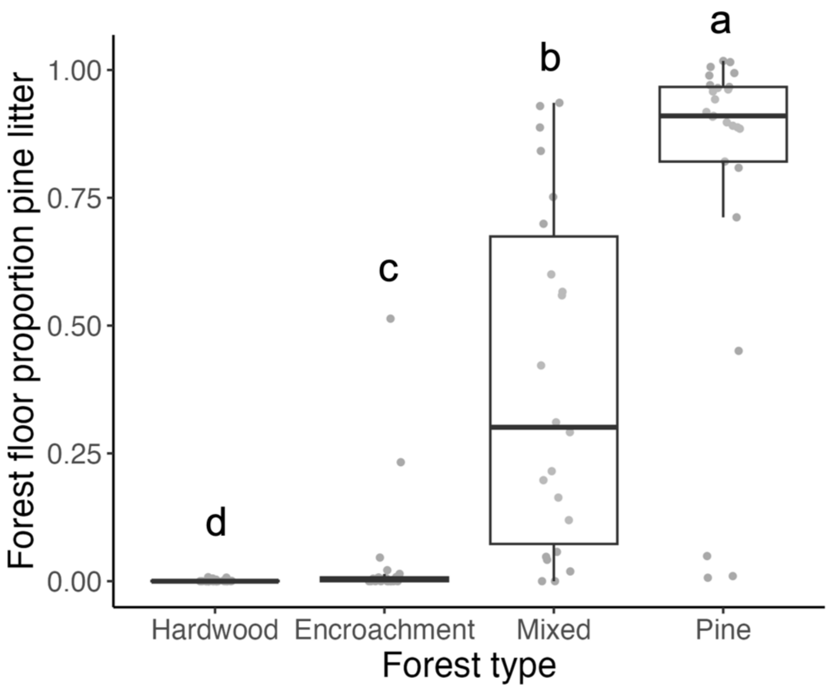


**Fig. S2.** Proportion of pine litter in the forest floor litter layer by forest type. Letters above each boxplot denote statistically significant differences between forest types as determined by pairwise Kruskal-Wallis tests.

**Table S1. Groups of Enzyme Commission (E.C.) numbers used to calculate relative abundances of genes encoding nutrient cycling enzymes**

| Enzyme type | E.C. number groups* | Relevant process(es) |
|---|---|---|
| Amidine hydrolases | 3.5.2, 3.5.3, 3.5.4 | Ammonification |
| Amino hydrolases | 3.5.5, 3.5.99 | Ammonification |
| Ammonium uptake | 1.14.18, 1.14.99.39, 6.3.1, 6.3.4 | Ammonification, nitrification |
| Chitinases | 3.2.1.14, 3.2.1.200-201 | Ammonification |
| Glycosidases, hydrolysing N-glycosyl compounds | 3.2.2 | Ammonification |
| Glycosidases, hydrolysing O- and S-glycosyl compounds | 3.2.1.50-184 | Ammonification |
| Oxidoreductases | 1.1.1, 1.1.2, 1.1.3, 1.1.5, 1.1.9, 1.1.98, 1.1.99, 1.10.1, 1.10.3, 1.10.5, 1.10.9, 1.10.99, 1.11.1, 1.11.2, 1.12.7, 1.12.98, 1.13.11, 1.13.12, 1.13.99, 1.14.1, 1.14.11, 1.14.12, 1.14.13, 1.14.14, 1.14.15, 1.14.16, 1.14.17, 1.14.18, 1.14.19, 1.14.20, 1.14.3, 1.14.99, 1.16.99, 1.17.1, 1.17.2, 1.17.3, 1.17.4, 1.17.5, 1.17.7, 1.17.98, 1.17.99, 1.2.1, 1.2.3, 1.2.7, 1.2.99, 1.20.1, 1.21.3, 1.21.4, 1.21.98, 1.3.1, 1.3.3, 1.3.5, 1.3.7, 1.3.8, 1.3.98, 1.3.99, 1.4.1, 1.4.2, 1.4.3, 1.4.4, 1.4.5, 1.4.7, 1.4.9, 1.4.98, 1.4.99, 1.5.1, 1.5.3, 1.5.4, 1.5.5, 1.5.7, 1.5.8, 1.5.98, 1.5.99, 1.6.4, 1.6.6, 1.6.99, 1.7.1, 1.7.2, 1.7.3, 1.7.99, 1.8.1, 1.8.2, 1.8.3, 1.8.4, 1.8.5, 1.8.6, 1.8.7, 1.8.98, 1.8.99, 1.99.1 | Ammonification, phosphate change |
| Peptidases | 3.4.1, 3.4.11, 3.4.12, 3.4.13, 3.4.14, 3.4.15, 3.4.16, 3.4.17, 3.4.18, 3.4.19, 3.4.2, 3.4.21, 3.4.22, 3.4.23, 3.4.24, 3.4.25, 3.4.3, 3.4.4, 3.4.99 | Ammonification |
| Phosphatases | 3.6.1, 3.9.1 | Phosphate change |
| Diphosphoric monoester hyrdolases | 3.1.7 | Phosphate change |
| Hydrolases acting on carbon-phosphorus bonds | 3.11.1 | Phosphate change |
| P-cycling hydrolases (Phosphoric monoester, diester, and triester, and triphosphoric monoester) | 3.1.3, 3.1.4, 3.1.5, 3.1.8 | Phosphate change |
| Nitrate reductase | 1.7.5 | Nitrification |
| Ammonia monooxygenase | 1.14.18, 1.14.99.39 | Nitrification |

*The numbers listed here represent groups of E.C. numbers (e.g., 1.1.1 contains enzymes 1.1.1.1, 1.1.1.2, etc.) for conciseness. Within the E.C. groups listed, only E.C. numbers encoding the enzyme type listed in column 1 were included in analysis.

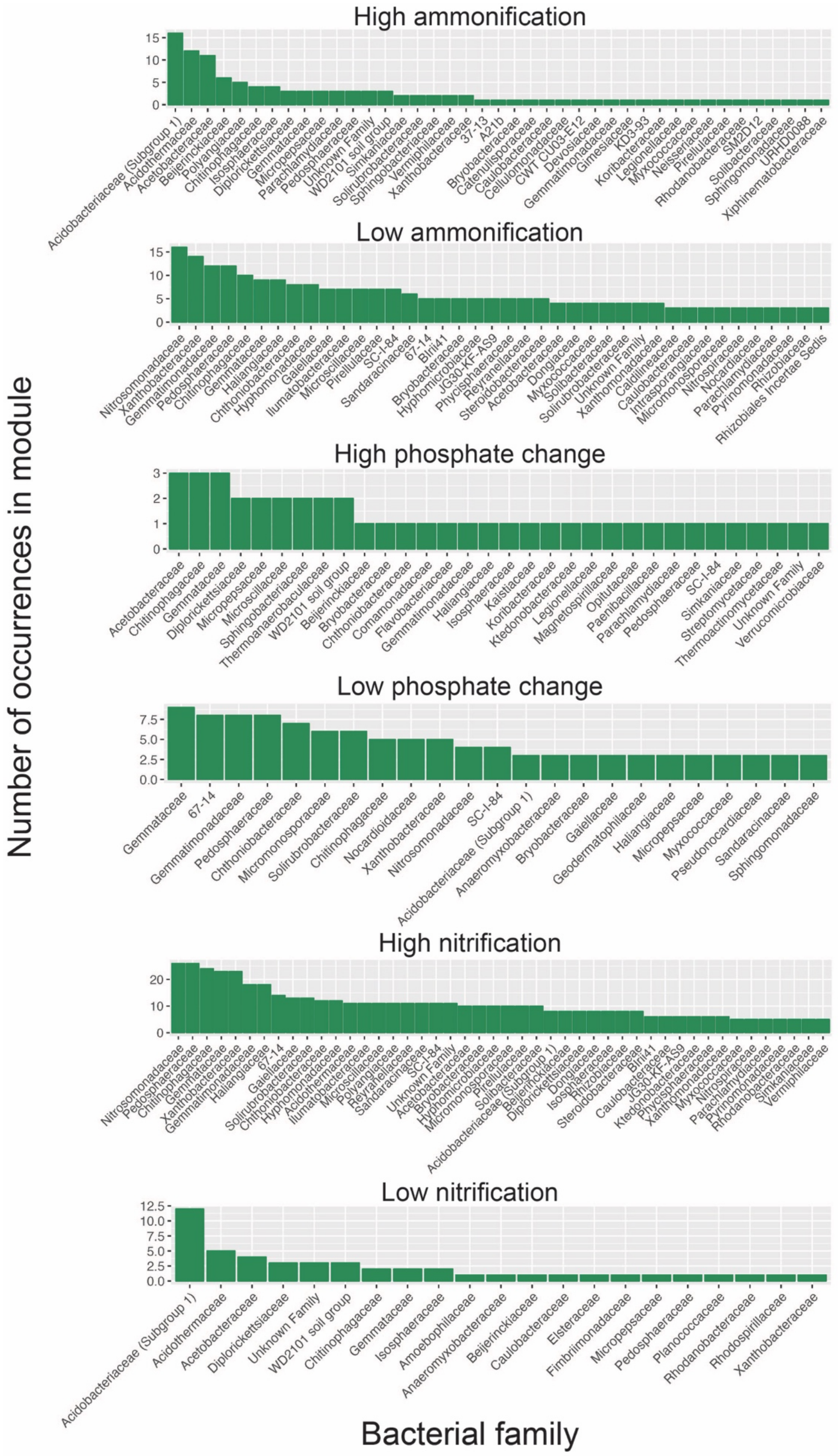

High ammonification
Low ammonification
High phosphate change
Low phosphate change
High nitrification
Low nitrification
Number of occurrences in module
Bacterial family

**Fig. S3.** Number of bacterial ASVs grouped by family in the bacterial indicator co-occurrence modules. The relative abundances of each ASV in the module were summed in each sample to calculate the relative abundance of the indicator co-occurrence module per sample. For the low ammonification and low phosphate change modules, which had high numbers of taxa, only bacterial families with more than 2 ASVs in the module are shown, and the high nitrification module only shows families with more than 4 ASVs.

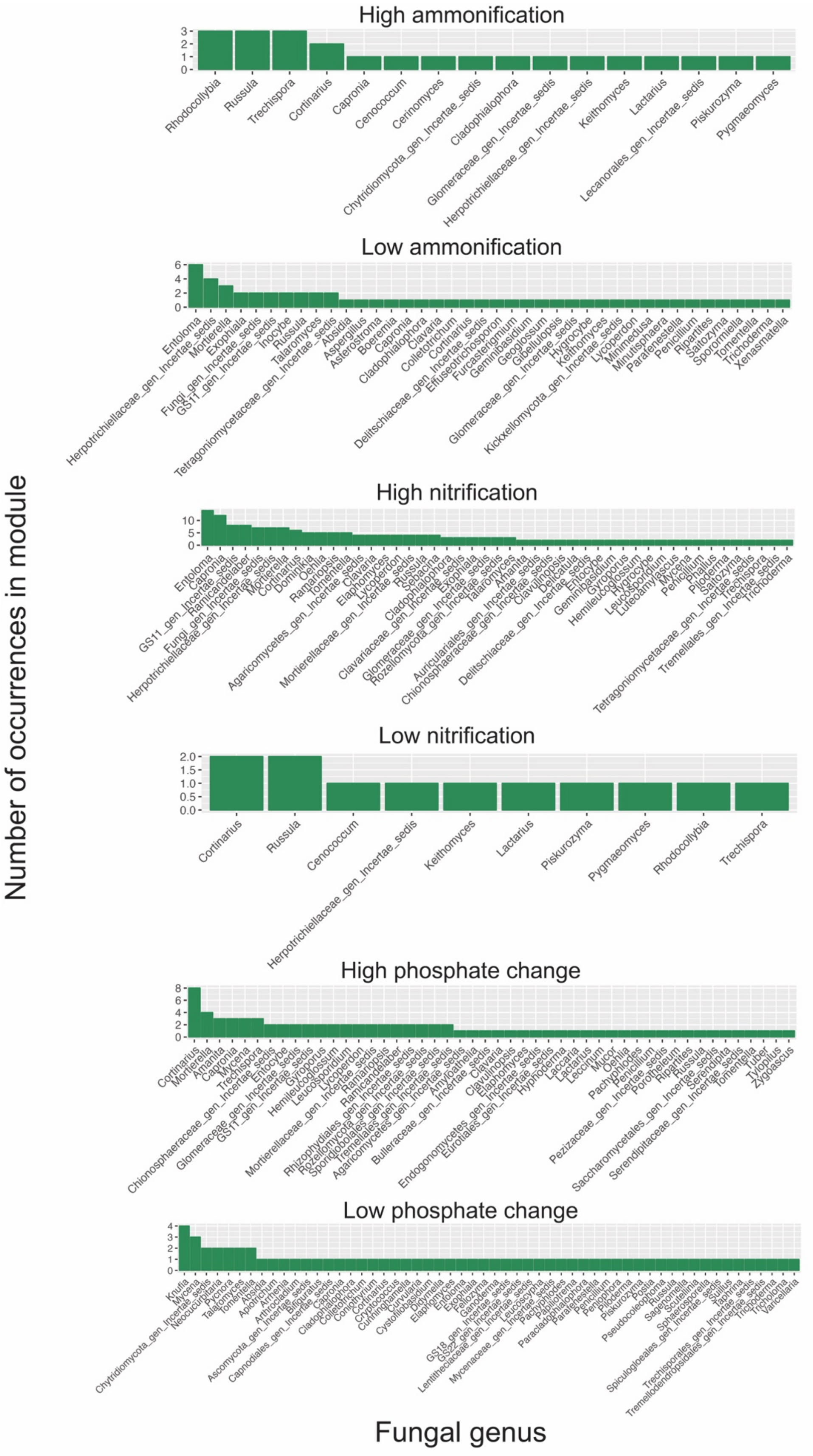

High ammonification
Rhodocollybia
Russula
Trechispora
Cortinarius
Capronia
Cenococcum
Cerinomyces
Chytridiomycota_gen_Incertae_sedis
Cladophialophora
Glomeraceae_gen_Incertae_sedis
Herpotrichiellaceae_gen_Incertae_sedis
Keithomyces
Lactarius
Lecanorales_gen_Incertae_sedis
Piskurozyma
Pygmaeomyces
Low ammonification
High nitrification
Low nitrification
Cortinarius
Russula
Cenococcum
Herpotrichiellaceae_gen_Incertae_sedis
Keithomyces
Lactarius
Piskurozyma
Pygmaeomyces
Rhodocollybia
Trechispora
High phosphate change
Low phosphate change
Number of occurrences in module
Fungal genus

**Fig. S4.** Number of fungal ASVs grouped by genus in the fungal indicator co-occurrence modules. The relative abundances of each ASV in the module were summed in each sample to calculate the relative abundance of the indicator co-occurrence module per sample. The fungal high nitrification module had a high number of genera, so only genera with more than 1 ASV in the module are shown.

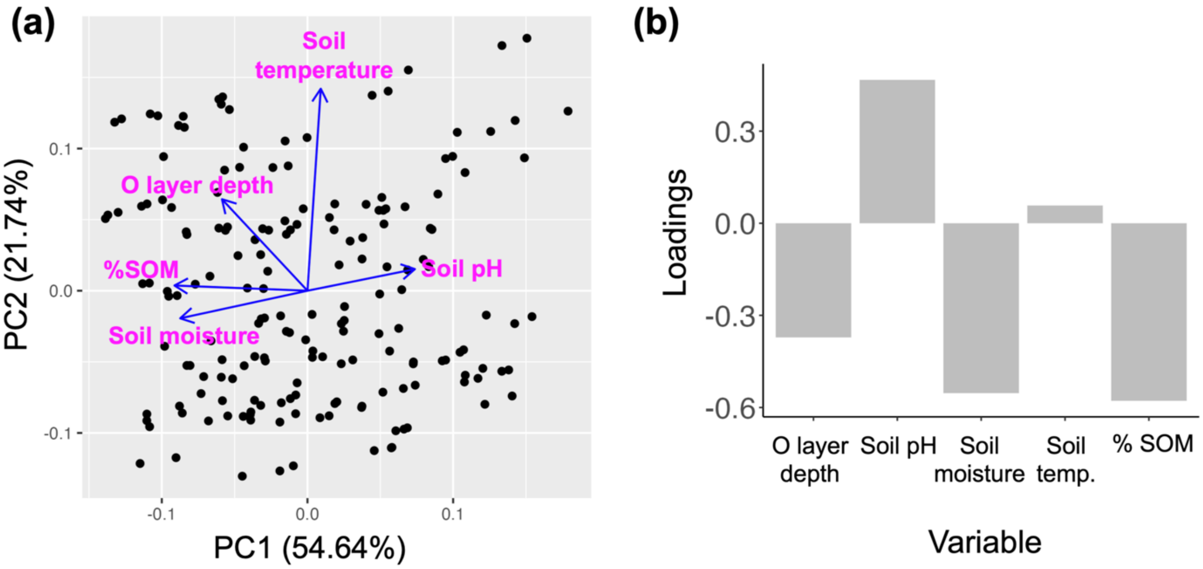


**Fig. S5.** Principal Component Analysis of soil edaphic variables. **(a)** Variables are plotted in principal component (PC) space. The percent of variation in the data explained by the first and second PC axes are shown in parentheses in each axis title. **(b)** The loadings for each variable in the first PC axis, which was used to explain soil nutrient cycling rates.

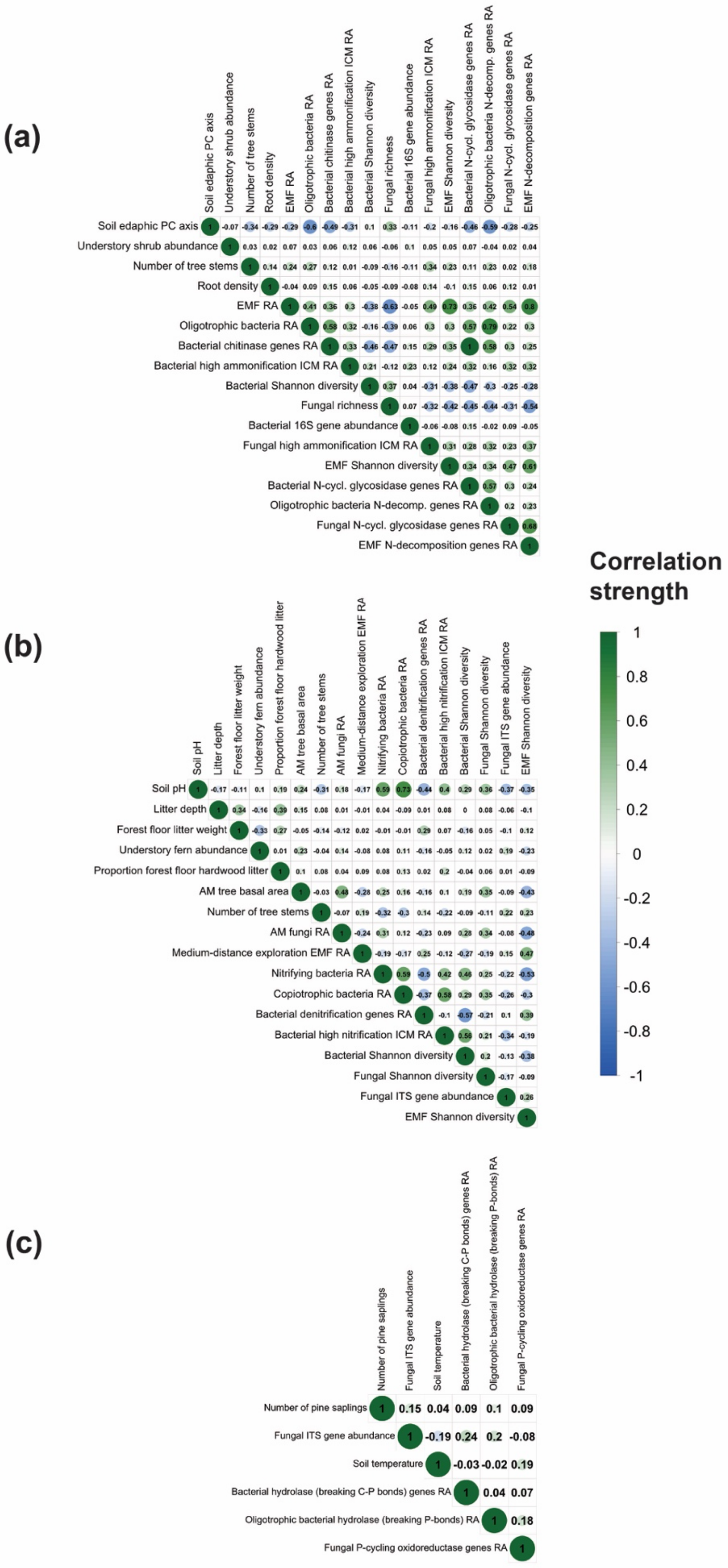
(a)
Soil edaphic PC axis
Understory shrub abundance
Number of tree stems
Root density
EMF RA
Oligotrophic bacteria RA
Bacterial chitinase genes RA
Bacterial high ammonification ICM RA
Bacterial Shannon diversity
Fungal richness
Bacterial 16S gene abundance
Fungal high ammonification ICM RA
EMF Shannon diversity
Bacterial N-cycl. glycosidase genes RA
Oligotrophic bacteria N-decomp. genes RA
Fungal N-cycl. glycosidase genes RA
EMF N-decomposition genes RA
Correlation strength
1
0.8
0.6
0.4
0.2
0
-0.2
-0.4
-0.6
-0.8
-1
(b)
Soil pH
Litter depth
Forest floor litter weight
Understory fern abundance
Proportion forest floor hardwood litter
AM tree basal area
Number of tree stems
AM fungi RA
Medium-distance exploration EMF RA
Nitrifying bacteria RA
Copiotrophic bacteria RA
Bacterial denitrification genes RA
Bacterial high nitrification ICM RA
Bacterial Shannon diversity
Fungal Shannon diversity
Fungal ITS gene abundance
EMF Shannon diversity
(c)
Number of pine saplings
Fungal ITS gene abundance
Soil temperature
Bacterial hydrolase (breaking C-P bonds) genes RA
Oligotrophic bacterial hydrolase (breaking P-bonds) RA
Fungal P-cycling oxidoreductase genes RA

**Fig. S6.** Pearson correlations between all explanatory variables input into the boosted regression tree modeling and multivariate linear model selection process for **(a)** net ammonification, **(b)** net nitrification, and **(c)** net phosphate change. The only exception was for the multivariate linear model selection for net nitrification, where input variables were reduced further as described in Appendix S1: Section S1: Methods and Appendix S1: Fig. S7. Variable name abbreviations include relative abundance (RA), ectomycorrhizal fungi (EMF), arbuscular mycorrhizal (AM), and indicator co-occurrence module (ICM).

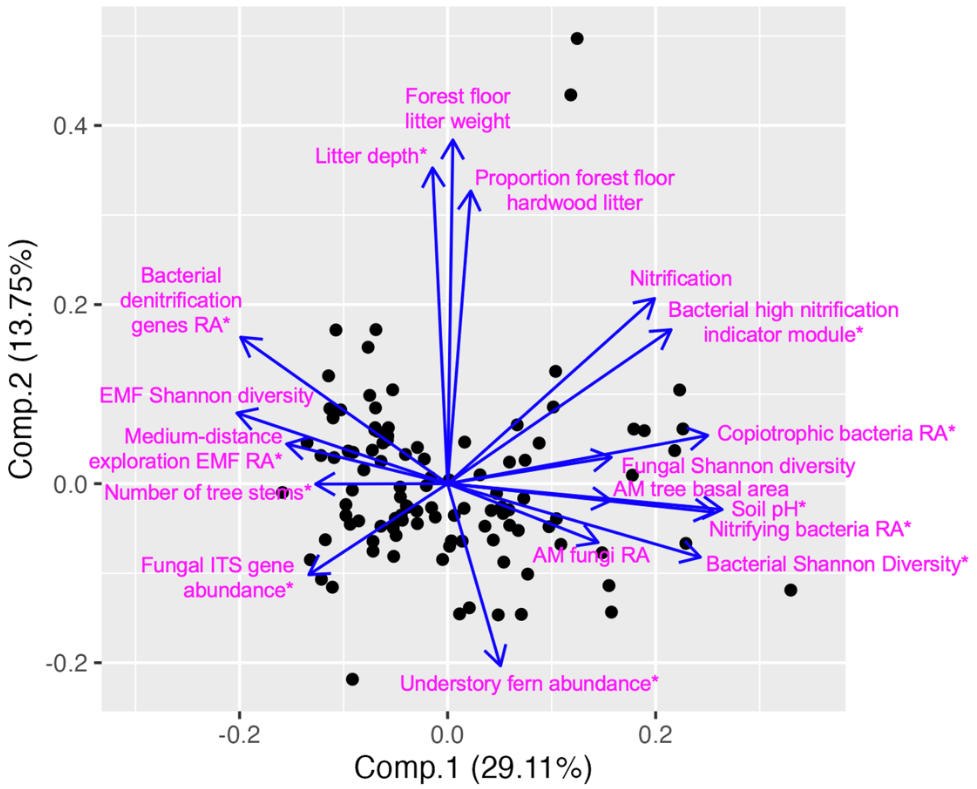


**Fig S7.** Principal Component Analysis (PCA) of all variables included in the Boosted Regression Tree analysis explaining net nitrification. When variables were highly correlated in PCA space, only the one with the strongest correlation with net nitrification was chosen for the multivariate linear model selection process. Variables input into the linear model selection process are denoted with a star (*). In variable names, relative abundance is abbreviated as RA.

<u>SECTION S2: RESULTS</u>

| Category | Variable | Correlation | P value |
|---|---|---|---|
| Microbial: functional guild relative abundances | Ectomycorrhizal fungi | 0.29 | 0.002 |
| | Saprotrophic fungi | -0.24 | 0.01 |
| | Arbuscular mycorrhizal fungi | -0.10 | 0.29 |
| | Fungal root endophyte fungi | -0.12 | 0.21 |
| | EMF contact exploration type | 0.07 | 0.48 |
| | EMF short-distance exploration type | -0.02 | 0.83 |
| | EMF medium-distance exploration type | 0.24 | 0.01 |
| | EMF long-distance exploration type | 0.00 | 0.97 |
| | Copiotrophic bacteria | -0.27 | 0.003 |
| | Oligotrophic bacteria | 0.39 | <0.001 |
| Microbial: functional gene relative abundances | Bacterial chitinase genes | 0.43 | <0.001 |
| | Fungal chitinase genes | 0.16 | 0.10 |
| | Fungal ammonium uptake genes | 0.09 | 0.35 |
| | Bacterial amidine hydrolase genes | 0.33 | <0.001 |
| | Bacterial amino hyrolase genes | 0.23 | 0.01 |
| | Bacterial N-cycling glycosidase (hydrolysing N-Glycosyl Compounds) genes | 0.37 | <0.001 |
| | Bacterial N-cycling glycosidase (hydrolysing O- and S-Glycosyl Compounds) genes | 0.42 | <0.001 |
| | Bacterial N-cycling oxidoreductase genes | 0.32 | <0.001 |
| | Bacterial amine oxidase genes | 0.26 | 0.004 |
| | Bacterial peptidase genes | 0.36 | <0.001 |
| | Fungal amidine hydrolase genes | 0.18 | 0.07 |
| | Fungal amino hyrolase genes | 0.14 | 0.13 |
| | Fungal N-cycling glycosidase (hydrolysing N-Glycosyl Compounds) genes | 0.18 | 0.06 |
| | Fungal N-cycling glycosidase (hydrolysing O- and S-Glycosyl Compounds) genes | 0.19 | 0.05 |
| | Fungal peptidase genes | 0.13 | 0.16 |
| | Fungal N-cycling oxidoreductase genes | 0.15 | 0.12 |
| | Fungal amine oxidase genes | 0.04 | 0.70 |
| Microbial: functional gene relative abundances within functional guilds | Oligotrophic bacterial chitinase genes | 0.31 | <0.001 |
| | Copiotrophic bacterial amidine hydrolase genes | -0.41 | <0.001 |
| | Oligotrophic bacterial amidine hydrolase genes | 0.35 | <0.001 |
| | Copiotrophic bacterial amino hyrolase genes | -0.27 | 0.003 |
| | Oligotrophic bacterial amino hyrolase genes | 0.35 | <0.001 |
| | Copiotrophic bacterial N-cycling glycosidase (hydrolysing N-Glycosyl Compounds) genes | -0.41 | <0.001 |
| | Oligotrophic bacterial N-cycling glycosidase (hydrolysing N-Glycosyl Compounds) genes | 0.37 | <0.001 |
| | Copiotrophic bacterial N-cycling glycosidase (hydrolysing O- and S-Glycosyl Compounds) genes | -0.42 | <0.001 |
| | Oligotrophic bacterial N-cycling glycosidase (hydrolysing O- and S-Glycosyl Compounds) genes | 0.23 | 0.01 |
| | Copiotrophic bacterial N-cycling oxidoreductase genes | -0.43 | <0.001 |
| | Oligotrophic bacterial N-cycling oxidoreductase genes | 0.34 | <0.001 |
| | Copiotrophic bacterial amine oxidase genes | -0.39 | <0.001 |
| | Oligotrophic bacterial amine oxidase genes | 0.04 | 0.65 |
| | Copiotrophic bacterial peptidase genes | -0.44 | <0.001 |
| | Oligotrophic bacterial peptidase genes | 0.35 | <0.001 |
| | All oligotrophic bacterial N-decomposition genes | 0.36 | <0.001 |
| | EMF chitinase genes | 0.22 | 0.02 |
| | Saprotrophic chitinase genes | -0.18 | 0.06 |
| | EMF ammonium uptake genes | 0.15 | 0.13 |
| | Saprotrophic ammonium uptake genes | -0.08 | 0.40 |
| | EMF amidine hydrolase genes | 0.23 | 0.02 |
| | Saprotrophic amidine hydrolase genes | -0.17 | 0.07 |
| | EMF amino hyrolase genes | 0.25 | 0.007 |
| | Saprotrophic amino hyrolase genes | -0.22 | 0.02 |
| | EMF N-cycling glycosidase (hydrolysing N-Glycosyl Compounds) genes | 0.23 | 0.01 |
| | Saprotrophic N-cycling glycosidase (hydrolysing N-Glycosyl Compounds) genes | -0.18 | 0.06 |
| | EMF N-cycling glycosidase (hydrolysing O- and S-Glycosyl Compounds) genes | 0.23 | 0.01 |
| | Saprotrophic N-cycling glycosidase (hydrolysing O- and S-Glycosyl Compounds) genes | -0.17 | 0.07 |
| | EMF peptidase genes | 0.23 | 0.02 |
| | Saprotrophic peptidase genes | -0.17 | 0.07 |
| | EMF N-cycling oxidoreductase genes | 0.24 | 0.01 |
| | Saprotrophic N-cycling oxidoreductase genes | -0.16 | 0.08 |
| | EMF amine oxidase genes | 0.17 | 0.08 |
| | Saprotrophic amine oxidase genes | -0.13 | 0.17 |
| | All EMF N-decomposition genes | 0.24 | 0.01 |
| Microbial: indicator co-occurrence modules | Fungal low ammonification indicator module | 0.19 | 0.05 |
| | Bacterial high ammonification indicator module | 0.26 | 0.00 |
| | Bacterial low ammonification indicator module | -0.21 | 0.02 |
| | Fungal high ammonification indicator module | 0.33 | <0.001 |
| Microbial: diversity metrics | Bacterial richness | -0.27 | 0.003 |
| | Bacterial Shannon diversity | -0.27 | 0.002 |
| | Fungal richness | -0.25 | 0.008 |
| | Fungal Shannon diversity | -0.19 | 0.05 |
| | Bacterial evenness | -0.20 | 0.03 |
| | Fungal evenness | -0.14 | 0.16 |
| | EMF richness | 0.13 | 0.18 |
| | EMF Shannon diversity | 0.20 | 0.03 |
| Microbial: 16S/ITS gene abundances | Bacterial 16S gene abundance | 0.22 | 0.01 |
| | Fungal ITS gene abundance | 0.05 | 0.57 |
| Soil edaphic factors | Soil pH | -0.42 | <0.001 |
| | Soil temperature | 0.09 | 0.35 |
| | O layer depth | 0.24 | 0.01 |
| | % soil moisture | 0.41 | <0.001 |
| | % soil organic matter | 0.43 | <0.001 |
| | Soil % nitrogen | 0.48 | <0.001 |
| | Soil % carbon | 0.46 | <0.001 |
| | Soil edaphic PC axis | -0.50 | <0.001 |
| Vegetation factors | Litter depth | 0.14 | 0.11 |
| | Proportion forest floor hardwood litter | -0.06 | 0.49 |
| | Total forest floor litter weight | 0.14 | 0.12 |
| | Total forest floor pine litter weight | 0.05 | 0.59 |
| | Total forest floor hardwood litter weight | -0.02 | 0.79 |
| | Abundance of broad-leaf herbs in the understory | 0.00 | 0.98 |
| | Abundance of grassy herbs in the understory | -0.14 | 0.11 |
| | Abundance of ferns in the understory | -0.02 | 0.81 |
| | Abundance of shrubs in the understory | 0.16 | 0.08 |
| | Maximum height of understory vegitation | -0.08 | 0.36 |
| | Abundance of all understory plants | -0.04 | 0.62 |
| | Arbuscular mycorrhizal tree basal area | -0.06 | 0.50 |
| | Ectomycorrhizal tree basal area | -0.01 | 0.95 |
| | Hardwood tree basal area | 0.06 | 0.51 |
| | White pine basal area | -0.09 | 0.30 |
| | Number of tree stems > 5cm DBH | 0.20 | 0.03 |
| | Total basal area of all trees > 5cm DBH | -0.05 | 0.61 |
| | Ectomycorrhizal:arbuscular mycorrhizal tree basal area | 0.06 | 0.52 |
| | Number of white pine saplings | -0.12 | 0.17 |
| | Root density | 0.23 | 0.01 |

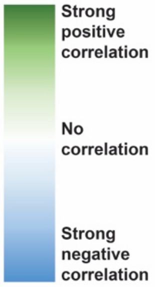

**Fig S8. Pearson correlations between net ammonification rates and all explanatory variables.** Statistically significant P-values < 0.05 are highlighted in yellow.

| | | Correlation | P value |
|---|---|---|---|
| Microbial: functional guild relative abundances | Ectomycorrhizal fungi | -0.37 | <0.001 |
| | Saprotrophic fungi | 0.28 | 0.002 |
| | Arbuscular mycorrhizal fungi | 0.29 | 0.002 |
| | Fungal root endophyte fungi | 0.05 | 0.59 |
| | EMF contact exploration type | -0.08 | 0.41 |
| | EMF short-distance exploration type | 0.08 | 0.37 |
| | EMF medium-distance exploration type | -0.32 | <0.001 |
| | EMF long-distance exploration type | -0.09 | 0.37 |
| | Nitrifying bacteria | 0.32 | <0.001 |
| | Copiotrophic bacteria | 0.48 | <0.001 |
| | Oligotrophic bacteria | -0.28 | 0.001 |
| Microbial: functional gene relative abundances | Bacterial denitrification genes | -0.24 | 0.008 |
| | Fungal denitrification genes | -0.03 | 0.78 |
| | Fungal ammonium uptake genes | -0.15 | 0.12 |
| Microbial: functional gene relative abundances within functional guilds | Saprotrophic fungal denitrification genes | 0.10 | 0.29 |
| | EMF ammonium uptake genes | -0.07 | 0.43 |
| | Saprotrophic fungal ammoinum uptake genes | 0.00 | 0.98 |
| | Nitrifying bacterial denitrification genes | 0.13 | 0.15 |
| | Copiotrophic bacterial denitrification genes | 0.24 | 0.006 |
| Microbial: indicator co-occurrence modules | Fungal high nitrification indicator module | -0.06 | 0.52 |
| | Fungal low nitrification indicator module | -0.17 | 0.07 |
| | Bacterial high nitrification indicator module | 0.44 | <0.001 |
| | Bacterial low nitrification indicator module | -0.07 | 0.41 |
| Microbial: diversity metrics | Bacterial richness | 0.36 | <0.001 |
| | Bacterial Shannon diversity | 0.37 | <0.001 |
| | Fungal richness | 0.32 | <0.001 |
| | Fungal Shannon diversity | 0.25 | 0.007 |
| | Bacterial evenness | 0.20 | 0.02 |
| | Fungal evenness | 0.18 | 0.06 |
| | EMF richness | -0.22 | 0.02 |
| | EMF Shannon diversity | -0.33 | <0.001 |
| Microbial: 16S/ITS gene abundances | Bacterial 16S gene abundance | 0.10 | 0.26 |
| | Fungal ITS gene abundance | -0.21 | 0.02 |
| Soil edaphic factors | Soil pH | 0.38 | <0.001 |
| | Soil temperature | 0.07 | 0.48 |
| | O layer depth | 0.02 | 0.82 |
| | % soil moisture | -0.08 | 0.35 |
| | % soil organic matter | -0.14 | 0.10 |
| | Soil % nitrogen | -0.06 | 0.50 |
| | Soil % carbon | -0.14 | 0.14 |
| | Soil edaphic PC axis | 0.22 | 0.02 |
| Vegetation factors | Litter depth | 0.28 | <0.001 |
| | Proportion forest floor hardwood litter | -0.09 | 0.31 |
| | Total forest floor litter weight | 0.23 | 0.009 |
| | Total forest floor pine litter weight | 0.03 | 0.78 |
| | Total forest floor hardwood litter weight | 0.27 | 0.002 |
| | Abundance of broad-leaf herbs in the understory | 0.07 | 0.43 |
| | Abundance of grassy herbs in the understory | 0.08 | 0.34 |
| | Abundance of ferns in the understory | -0.17 | 0.05 |
| | Abundance of shrubs in the understory | -0.03 | 0.76 |
| | Maximum height of understory vegitation | 0.11 | 0.21 |
| | Abundance of all understory plants | -0.03 | 0.75 |
| | Arbuscular mycorrhizal tree basal area | 0.35 | <0.001 |
| | Ectomycorrhizal tree basal area | -0.15 | 0.10 |
| | Hardwood tree basal area | 0.08 | 0.39 |
| | White pine basal area | 0.00 | 0.97 |
| | Number of tree stems > 5cm DBH | -0.18 | 0.03 |
| | Total basal area of all trees > 5cm DBH | 0.06 | 0.51 |
| | Ectomycorrhizal:arbuscular mycorrhizal tree basal area | -0.12 | 0.16 |
| | Number of white pine saplings | -0.14 | 0.12 |
| | Root density | -0.01 | 0.87 |

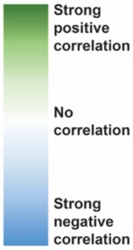


**Fig S9. Pearson correlations between net nitrification rates and all explanatory variables.**

Statistically significant P-values < 0.05 are highlighted in yellow.

| Category | Variable | Correlation | P value |
|---|---|---|---|
| Microbial: functional guild relative abundances | Ectomycorrhizal fungi | 0.05 | 0.61 |
| | Saprotrophic fungi | -0.04 | 0.68 |
| | Arbuscular mycorrhizal fungi | 0.03 | 0.78 |
| | Fungal root endophyte fungi | 0.07 | 0.51 |
| | EMF contact exploration type | -0.03 | 0.79 |
| | EMF short-distance exploration type | -0.15 | 0.14 |
| | EMF medium-distance exploration type | 0.13 | 0.20 |
| | EMF long-distance exploration type | 0.08 | 0.44 |
| | Copiotrophic bacteria | -0.02 | 0.82 |
| | Oligotrophic bacteria | 0.13 | 0.18 |
| Microbial: functional gene relative abundances | Fungal phosphate uptake genes | -0.02 | 0.85 |
| | Bacterial diphosphoric monoester hydrolase genes | -0.11 | 0.27 |
| | Bacterial hydrolase (acting on C-P bonds) genes | 0.26 | 0.007 |
| | Bacterial P-cycling oxidoreductase genes | 0.11 | 0.27 |
| | Bacterial phosphatase genes | 0.08 | 0.40 |
| | Bacterial all P-cycling hydrolase genes | 0.10 | 0.33 |
| | Fungal diphosphoric monoester hydrolase genes | 0.02 | 0.84 |
| | Fungal phosphatase genes | 0.10 | 0.33 |
| | Fungal all P-cycling hydrolase genes | 0.16 | 0.12 |
| | Fungal P-cycling oxidoreductase genes | 0.21 | 0.04 |
| Microbial: functional gene relative abundances withinfunctional guilds | EMF phosphate uptake genes | -0.04 | 0.67 |
| | Saprotrophic fungal phosphate uptake genes | 0.06 | 0.57 |
| | Copiotrophic bacterial diphosphoric monoester hydrolase genes | -0.07 | 0.45 |
| | Copiotrophic bacterial P-cycling oxidoreductase genes | -0.09 | 0.36 |
| | Oligotrophic bacterial P-cycling oxidoreductase genes | 0.12 | 0.22 |
| | Copiotrophic bacterial phosphatase genes | -0.03 | 0.72 |
| | Oligotrophic bacterial phosphatase genes | 0.14 | 0.14 |
| | Copiotrophic bacterial all P-cycling hydrolase genes | -0.08 | 0.39 |
| | Oligotrophic bacterial all P-cycling hydrolase genes | 0.16 | 0.10 |
| | EMF diphosphoric monoester hydrolase genes | 0.04 | 0.67 |
| | Saprotrophic fungal diphosphoric monoester hydrolase genes | -0.04 | 0.70 |
| | EMF phosphatase genes | 0.10 | 0.34 |
| | Saprotrophic fungal phosphatase genes | -0.07 | 0.48 |
| | EMF all P-cycling hydrolase genes | 0.10 | 0.32 |
| | Saprotrophic fungal all P-cycling hydrolase genes | -0.08 | 0.45 |
| | EMF P-cycling oxidoreductase genes | 0.09 | 0.38 |
| | Saprotrophic fungal P-cycling oxidoreductase genes | -0.06 | 0.55 |
| Microbial: indicator co-occurrence modules | Fungal high phosphate release indicator module | -0.03 | 0.76 |
| | Fungal low phosphate release indicator module | 0.03 | 0.80 |
| | Bacterial high phosphate release indicator module | 0.04 | 0.70 |
| | Bacterial low phosphate release indicator module | -0.14 | 0.16 |
| Microbial: diversity metrics | Bacterial richness | -0.04 | 0.66 |
| | Bacterial Shannon diversity | -0.03 | 0.75 |
| | Fungal richness | -0.03 | 0.78 |
| | Fungal Shannon diversity | 0.11 | 0.29 |
| | Bacterial evenness | -0.01 | 0.93 |
| | Fungal evenness | 0.15 | 0.15 |
| | EMF richness | 0.05 | 0.59 |
| | EMF Shannon diversity | 0.12 | 0.22 |
| Microbial: 16S/ITS gene abundances | Bacterial 16S gene abundance | -0.11 | 0.27 |
| | Fungal ITS gene abundance | 0.22 | 0.02 |
| Soil edaphic factors | Soil pH | -0.12 | 0.20 |
| | O layer depth | -0.08 | 0.41 |
| | % soil organic matter | 0.11 | 0.23 |
| | Soil % nitrogen | 0.08 | 0.43 |
| | Soil % carbon | 0.11 | 0.28 |
| | Soil edaphic PC axis | -0.11 | 0.27 |
| | Soil temperature | -0.21 | 0.02 |
| | % soil moisture | 0.08 | 0.40 |
| Vegetation factors | Litter depth | 0.04 | 0.70 |
| | Proportion forest floor hardwood litter | -0.11 | 0.27 |
| | Total forest floor litter weight | -0.15 | 0.11 |
| | Total forest floor pine litter weight | 0.04 | 0.66 |
| | Total forest floor hardwood litter weight | -0.12 | 0.23 |
| | Abundance of broad-leaf herbs in the understory | -0.09 | 0.32 |
| | Abundance of grassy herbs in the understory | 0.02 | 0.86 |
| | Abundance of ferns in the understory | 0.06 | 0.55 |
| | Abundance of shrubs in the understory | 0.06 | 0.51 |
| | Maximum height of understory vegitation | -0.01 | 0.94 |
| | Abundance of all understory plants | 0.11 | 0.26 |
| | Arbuscular mycorrhizal tree basal area | 0.14 | 0.14 |
| | Ectomycorrhizal tree basal area | -0.03 | 0.74 |
| | Hardwood tree basal area | 0.02 | 0.82 |
| | White pine basal area | 0.03 | 0.75 |
| | Number of tree stems > 5cm DBH | 0.06 | 0.53 |
| | Total basal area of all trees > 5cm DBH | 0.04 | 0.69 |
| | Ectomycorrhizal:arbuscular mycorrhizal tree basal area | -0.15 | 0.11 |
| | Number of white pine saplings | 0.22 | 0.02 |
| | Root density | -0.05 | 0.60 |

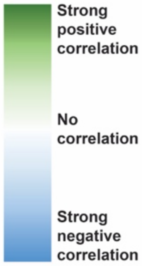


**Fig S10. Pearson correlations between net phosphate change rates and all explanatory variables.** Statistically significant P-values < 0.05 are highlighted in yellow.

**Table S2. Best-fit transect-level multivariate linear models explaining soil nutrient cycling rates based on minimum AIC value.**

| Response variable | Independent Variables | Coefficient | $R^2$ |
|---|---|---|---|
| Net ammonification | Soil temperature<br>% soil moisture<br>Medium-distance exploration type EMF relative abundance | 1.34$^{\dagger}$<br>57.87***<br>28.39** | 0.51 |
| Net soil phosphate change | Bacterial low phosphate change indicator module relative abundance<br>Litterfall magnesium concentration<br>Litterfall molybdenum concentration | -0.31<br>$-2.7e^{-05}$<br>0.04 | 0.47 |
| Net nitrification (log) | Copiotrophic bacteria relative abundance<br>Fungal Shannon diversity | 26.83**<br>1.19* | 0.55 |

Variable significance within model: † = $p < 0.1$, * = $p < 0.05$, ** = $p < 0.01$, *** = $p < 0.001$

Hartig, Florian. 2022. "DHARMa: Residual Diagnostics for Hierarchical (Multi-Level / Mixed) Regression Models."

Langfelder, Peter, and Steve Horvath. 2008. "WGCNA: An R Package for Weighted Correlation Network Analysis." *BMC Bioinformatics* 9 (1): 559. https://doi.org/10.1186/1471-2105-9-559.

Langfelder, Peter, and Steve Horvath. 2012. "Fast R Functions for Robust Correlations and Hierarchical Clustering." *Journal of Statistical Software* 46 (March):1–17.

Leathwick, Jr, J Elith, Mp Francis, T Hastie, and P Taylor. 2006. "Variation in Demersal Fish Species Richness in the Oceans Surrounding New Zealand: An Analysis Using Boosted Regression Trees." *Marine Ecology Progress Series* 321 (September):267–81.

Martin, Marcel. 2011. "Cutadapt Removes Adapter Sequences from High-Throughput Sequencing Reads." *EMBnet.Journal* 17 (1): 10–12.

Matteucci, Silvia, and Aída Colma. 1982. *Metodología Para El Estudio de La Vegetación / Por Silvia D. Matteucci y Aída Colma. SERBIULA (Sistema Librum 2.0).*

Nilsson, Rolf Henrik, Karl-Henrik Larsson, Andy F S Taylor, Johan Bengtsson-Palme, Thomas S Jeppesen, Dmitry Schigel, Peter Kennedy, et al. 2019. "The UNITE Database for Molecular Identification of Fungi: Handling Dark Taxa and Parallel Taxonomic Classifications." *Nucleic Acids Research* 47 (D1): D259–64.

Olsen, Sterling Robertson. 1954. *Estimation of Available Phosphorus in Soils by Extraction with Sodium Bicarbonate*. U.S. Department of Agriculture.

Quast, Christian, Elmar Pruesse, Pelin Yilmaz, Jan Gerken, Timmy Schweer, Pablo Yarza, Jörg Peplies, and Frank Oliver Glöckner. 2013. "The SILVA Ribosomal RNA Gene Database Project: Improved Data Processing and Web-Based Tools." *Nucleic Acids Research* 41 (Database issue): D590–96.

Wang, Qiong, George M. Garrity, James M. Tiedje, and James R. Cole. 2007. "Naive Bayesian Classifier for Rapid Assignment of rRNA Sequences into the New Bacterial Taxonomy." *Applied and Environmental Microbiology* 73 (16): 5261–67.
